# A JNK-interacting protein 1 acts across the midline to mediate synaptic localization of the SARM1 calcium-signaling scaffold protein for asymmetric neuronal fate choice

**DOI:** 10.64898/2026.04.30.722091

**Authors:** Yi-Wen Hsieh, Shengyao Yuan, Jun Yang, Cesar Siete, Chiou-Fen Chuang

## Abstract

The *Caenorhabditis elegans* AWC olfactory neuron pair specifies asymmetric subtypes, AWC^OFF^ and AWC^ON^, through stochastic and coordinated cell signaling events. UNC-104/kinesin-3 (KIF1A) and UNC-116/kinesin-1 motor proteins act in the AWC^ON^ cell to regulate the synaptic localization of the TIR-1/SARM1-assembled calcium signaling complex in the AWC^OFF^ cell to promote AWC^OFF^. However, the molecular mechanism in the AWC^ON^ cell that acts non-cell autonomously to control synaptic TIR-1 calcium signaling to promote AWC^OFF^ remains unclear. Here, we show that JIP-1, a conserved c-Jun N-terminal kinase (JNK)-interacting protein 1, mediates the synaptic localization of TIR-1 in the AWC axon to specify the AWC^OFF^ subtype. A *jip-1* loss-of-function mutant, identified from an unbiased forward genetic screen, has reduced localization of TIR-1 at synapses in the AWC axon and accumulation of TIR-1 in the AWC cell body. *jip-1* mutants significantly enhance the 2AWC^ON^ phenotype of a hypomorphic *tir-1* mutant. JIP-1, like UNC-104 and UNC-116, mainly acts non-cell autonomously in AWC^ON^ to specify the AWC^OFF^ subtype. Our findings provide mechanistic insights into how cell-specific Ca^2+^ signaling proteins, such as TIR-1, target synaptic regions via intercellular signaling to promote neuronal diversification.

## Introduction

The *C. elegans* left and right AWC olfactory neurons specify asymmetric subtypes, default AWC^OFF^ and induced AWC^ON^, with distinct functions through a stochastic and coordinated cell signaling event (TROEMEL *et al*. 1997; TAYLOR *et al*. 2010; ALQADAH *et al*. 2013; HSIEH *et al*. 2014; ALQADAH *et al*. 2016b; HSIEH *et al*. 2017; ALQADAH *et al*. 2018). Wild-type animals have one AWC^ON^ subtype, expressing the G protein-coupled receptor (GPCR) gene *str-2*, and one AWC^OFF^ subtype, expressing the GPCR gene *srsx-3* (TROEMEL *et al*. 1999; BAUER HUANG *et al*. 2007). AWC olfactory neuron subtypes are specified during embryogenesis and maintained throughout life (TROEMEL *et al*. 1999; CHUANG AND BARGMANN 2005; LESCH *et al*. 2009; LESCH AND BARGMANN 2010).

The AWC^OFF^ subtype is established using a calcium-activated MAP kinase cascade. The AWC^ON^ subtype is induced by the repression of the calcium signaling in the contralateral neuron via NSY-5 gap-junction-mediated intercellular signaling and SLO BK potassium channels (TROEMEL *et al*. 1999; SAGASTI *et al*. 2001; TANAKA-HINO *et al*. 2002; CHUANG AND BARGMANN 2005; VANHOVEN *et al*. 2006; BAUER HUANG *et al*. 2007; CHUANG *et al*. 2007; CHANG *et al*. 2011; HSIEH *et al*. 2012; SCHUMACHER *et al*. 2012; COCHELLA *et al*. 2014; ALQADAH *et al*. 2015; PAGANO *et al*. 2015; ALQADAH *et al*. 2016a; ALQADAH *et al*. 2019). TIR- 1/SARM1 acts cell autonomously, downstream of the initial intercellular signaling that establishes AWC asymmetry, to execute the AWC^OFF^ subtype (CHUANG AND BARGMANN 2005). TIR-1 functions as a scaffold protein to assemble a Ca^2+^-regulated signaling complex, consisting of UNC-43 calcium/calmodulin-dependent protein kinase (CaMKII) and NSY-1 MAP kinase kinase kinase (ASK1 MAPKKK), at postsynaptic regions in the AWC axon to specify AWC^OFF^ (TROEMEL *et al*. 1997; SAGASTI *et al*. 2001; CHUANG AND BARGMANN 2005; CHANG *et al*. 2011). UNC-104/kinesin-3 (KIF1A) and UNC-116/kinesin-1 motor proteins act non-cell autonomously to regulate the synaptic localization of the TIR-1 calcium-signaling complex to promote the AWC^OFF^ subtype (CHANG *et al*. 2011; KHALID *et al*. 2026). However, the molecular mechanism in the AWC^ON^ cell that mediates non-cell-autonomous control of synaptic TIR-1 calcium signaling to promote AWC^OFF^ remains to be determined.

Here, we identify the role of the JNK-interacting protein JIP-1 in the synaptic localization of TIR-1, thereby promoting the AWC^OFF^ subtype, from an unbiased forward genetic screen. Like the previously identified roles of UNC-104/kinesin-3 (KIF1A) and UNC-116/kinesin-1 in AWC asymmetry, JIP-1 mainly acts non-cell autonomously to specify AWC^OFF^. Our results suggest a model in which JIP-1 may function with UNC-104/kinesin-3 (KIF1A) and UNC-116/kinesin-1 to non-cell autonomously regulate the synaptic localization of the TIR-1 signaling complex in AWC^OFF^ for promoting its subtype.

## Results

### The *vy6* mutation causes a defect in the synaptic localization of TIR-1 required for promoting AWC^OFF^

The subcellular localization of TIR-1 in AWC neurons was examined using an integrated *odr-3p::tir-1::GFP* transgene, expressing functional TIR-1::GFP from an AWC *odr-3* promoter. In the wild type, TIR-1::GFP was localized in a punctate pattern along the AWC axon and primarily excluded from the cell body and dendrites, as previously described (CHUANG AND BARGMANN 2005; CHANG *et al*. 2011) (Figure 1A). Our previous study showed that UNC-104/kinesin-3 (KIF1A) and UNC-116/kinesin-1 motor proteins regulate the synaptic localization of the TIR-1 calcium-signaling complex in the AWC axon to specify the AWC^OFF^ subtype (CHANG *et al*. 2011; KHALID *et al*. 2026).

**Figure 1.**
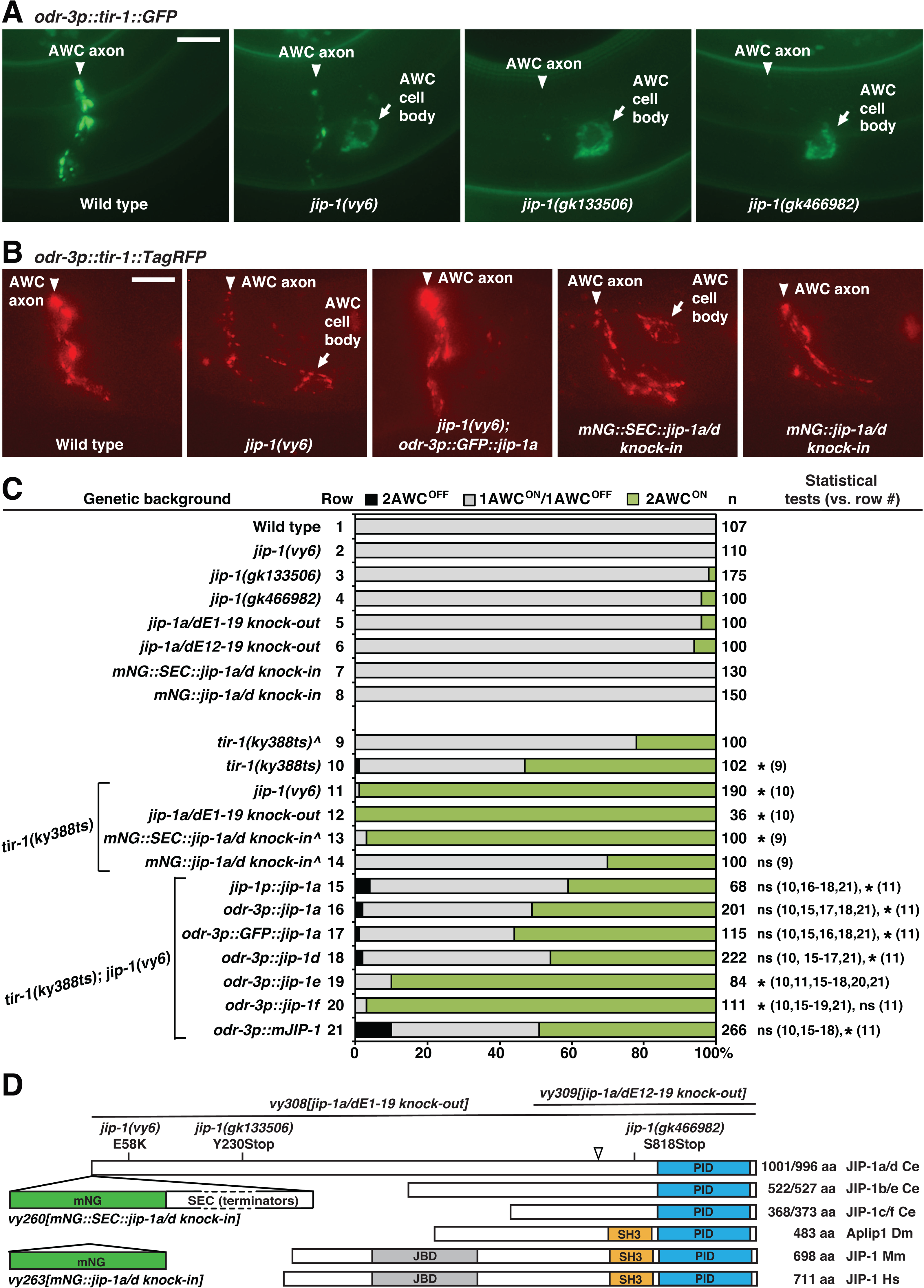
*jip-1* is required for synaptic localization of TIR-1 to promote AWC^OFF^. (**A**) Images of wild type and *jip-1* mutants expressing TIR-1::GFP in AWC cells from a stably integrated transgene *odr-3p::tir-1::GFP* at the first larval (L1) stage. In the wild type, TIR-1::GFP was localized in a punctate pattern along the AWC axon and primarily excluded from the cell body and dendrites. In *jip-1(vy6)*, *jip-1(gk133506)*, and *jip-1(gk466982)* mutants, the localization of TIR-1::GFP was reduced in the AWC axon and accumulated in the AWC cell body. The anterior is left, and the ventral is down. Scale bar, 5 μm. Quantification of TIR-1::GFP fluorescence intensity in the AWC axon and cell body is included in Figure S1A and S1B, respectively. (**B**) Images of TIR-1::TagRFP expression in AWC cells from a single copy insertion transgene *odr-3p::tir-1::TagRFP* at the L1 stage. The single copy insertion transgene *odr-3p::GFP::jip-1a* rescued the mislocalized TIR-1 phenotype in *jip-1(vy6)* mutants. *mNG::SEC::jip-1a/d knock-in* animals showed reduced TIR-1 localization in the AWC axon and accumulation of TIR-1 in the AWC cell body, similar to the phenotype observed in *jip-1* mutants*. mNG::jip-1a/d knock-in,* with SEC excised, had a wild-type TIR-1 localization pattern. The anterior is left, and the ventral is down. Scale bar, 5 μm. (**C**) Quantification of AWC^ON^ marker transgene, *str-2p::GFP*, expression in adults. Animals were grown at 20°C or 15°C (^). n, total number of animals scored. 2AWC^OFF^, *str-2* is not expressed in either AWC; 1AWC^OFF^/AWC^ON^, only one of the two AWC cells expresses *str-2;* 2AWC^ON^, *str-2* expressed in both AWC neurons. Statistical comparisons of 2AWC^ON^ phenotype penetrance were made using the *Z*-test. Asterisks indicate comparisons that are different at *p* < 0.05. ns, not significant. The structure of *jip-1* genomic DNA and transgenes are illustrated in Figure S2A. (**D**) Structure of JIP-1 proteins in *C. elegans* and other species. JBD, JNK-binding domain; SH3, Src homology 3 domain; PID, phosphotyrosine-interaction domain. Ce: *Caenorhabditis elegans*; Dm, *Drosophila melanogaster*; Mm, *Mus musculus*; Hs, *Homo sapiens*. *mNG*, mNeonGreen. SEC, self-excising cassette containing transcriptional terminators, a dominant roller phenotype marker *sqt-1(e1350),* Cre driven by a heat shock promoter, and a hygromycin resistance gene. The SEC cassette is flanked by LoxP sites. The arrowhead indicates the position at which the amino acid sequence differs between *C. elegans* JIP-1 protein isoforms a/e/f (SFFSPD) and b/c/d (Y). Amino acid sequence alignment of the PID domain in JIP-1 proteins is included in Figure S2B.

To identify additional molecules required for the synaptic localization of TIR-1 in AWC subtype choice, we performed an unbiased forward genetic screen to isolate mutants with defective TIR-1::GFP localization in AWC neurons. From the screen, we identified the *vy6* allele that displayed reduced localization of TIR-1::GFP at synapses in the AWC axon and accumulation of TIR-1::GFP in the AWC cell body (Figures 1A, S1A, and S1B). TIR-1::TagRFP, expressed from a single copy insertion transgene *odr-3p::tir-1::TagRFP*, displayed a localization pattern similar to TIR-1::GFP, expressed from a multiple copy insertion transgene *odr-3p::tir-1::GFP*, in wild type and *jip-1(vy6)* mutants (Figure 1A and 1B). Like *vy6* mutants, *unc-104(e1265)* and *unc-116(e2310)* mutants displayed reduced localization of TIR-1::GFP in the AWC axon, but TIR-1::GFP was not detected in the AWC cell body or dendrites in *unc-104(e1265)* or *unc-116(e2310)* mutants (CHANG *et al*. 2011; KHALID *et al*. 2026) (Figure S1C).

As a control, the fluorescence intensity of GFP in AWC cells was compared between wild type and *vy6* mutants containing the transgene *odr-3p::GFP*. The fluorescence intensity of GFP in the AWC cell body was not significantly different between wild type and *vy6* mutants (Figure S1D). This result rules out the possibility that the *odr-3* promoter activity is affected by the *vy6* mutation and supports the notion that the effect of the *vy6* mutation on TIR-1::GFP is at the subcellular localization level.

Similar to *unc-104(e1265)* mutants, *vy6* mutants alone did not show a defect in AWC asymmetry but significantly enhanced the penetrance of the 2AWC^ON^ phenotype (both AWC become AWC^ON^) of *tir-1(ky388)* temperature-sensitive (ts) mutants from 54% to 99% at 20°C (CHANG *et al*. 2011) (Figure 1C, rows 2, 10, and 11). These results suggest that the *vy6* mutation reduces synaptic localization of TIR-1 in the AWC axon, thereby increasing the penetrance of the 2AWC^ON^ phenotype in a hypomorphic *tir-1* mutant background.

### *vy6* is a missense mutation in the JNK-interacting protein JIP-1

We identified the molecular lesion in *vy6* mutants using one-step whole genome sequencing and single-nucleotide polymorphism mapping (DOITSIDOU *et al*. 2010). The *vy6* lesion was identified as a G to A mutation, resulting in a glutamic acid to lysine change in the second exon of F56D12.4a/d (*jip-1a/d*) isoforms (Figures 1D and S2A). *jip-1* has six alternatively spliced isoforms, a-f (wormbase.org) (Figures 1D and S2A). These isoforms are classified into three subtypes (a/d, b/e, and c/f), each including two isoforms of similar length with alternative transcription start, termination, or both sites. The coding regions of the two isoforms in each subtype differ only in the presence or absence of 15 base pairs at the beginning of an exon (exon 13 in *jip-1a/d*, exon 5 in *jip-1b/e*, and exon 3 in *jip-1c/f*), resulting in serine-phenylalanine-phenylalanine-serine-proline-aspartic acid (SFFSPD) sequence in JIP-1a/e/f and tyrosine (Y) in JIP-1d/b/c proteins (wormbase.org) (Figures 1D and S2A). The glutamic acid residue affected by the *vy6* mutation is present in JIP-1a/d but absent in JIP-1b/c/e/f (Figure 1D and S2A).

*jip-1* encodes a JNK-interacting protein JIP-1, an ortholog of human and mouse mitogen-activated protein kinase 8 interacting protein 1 (MAPK8IP1; also known as JIP-1 or JIP1) (wormbase.org; Mouse Genome Informatics; uniport.org). JIPs were first identified as a regulator of stress-induced JNK (DICKENS *et al*. 1997). JIPs are a family of scaffold proteins that localize pathway components to specific subcellular sites, facilitate their activation, and enable signal integration (WHITMARSH 2006). JIPs are highly conserved between *C. elegans, Drosophila*, and mammals. There are two predicted *jip* genes in *C. elegans* (wormbase.org), two in *Drosophila melanogaster* (flybase.org), four in mice (Mouse Genome Informatics), and four in humans (WHITMARSH 2006; DHANASEKARAN *et al*. 2007). Like other JIP-1 proteins, the *C. elegans* JIP-1 protein has a predicted phosphotyrosine-interaction domain (PID, phosphotyrosine-binding domain, or PTB) at the C-terminus (WHITMARSH 2006; DHANASEKARAN *et al*. 2007) (Figures 1D and S2A). The predicted PID domain of *C. elegans* JIP-1 and mammalian (mouse and human) JIP-1s are 42% identical and 75-88% similar (Figure S2B).

Two nonsense alleles of *jip-1*, *jip-1(gk133506)* and *jip-1(gk466982)*, were created through the million-mutation project (THOMPSON *et al*. 2013) (Figures 1D and S2A). The *jip-1(gk133506)* mutation results in a premature stop codon in the fifth exon of *jip-1a/d* but does not affect the coding region of *jip-1b,c,e,f*. The *jip-1(gk466982)* mutation causes a premature stop codon in the last fifth exon of all six *jip-1* isoforms. Both *jip-1(gk133506)* and *jip-1(gk466982)* mutants display reduced localization of TIR-1::GFP in the AWC axon and accumulation of TIR-1::GFP in the AWC cell body, similar to the phenotype observed in *jip-1(vy6)* mutants (Figures 1A, S1A, and S1B). These results support that the molecular lesion of *vy6* in *jip-1* causes the mislocalized TIR-1 phenotype.

We generated two *jip-1* deletion alleles, *jip-1a/dE1-19 knock-out* and *jip-1a/dE12-19 knock-out*, by replacing the entire coding sequence or the coding sequence from exons 12-19 of *jip-1a/d*, respectively, with the fluorescent reporter mNeonGreen (mNG) using Cas9-triggered homologous recombination (DICKINSON *et al*. 2013) (Figures 1D and S2A). Like *jip-1(gk133506)* and *jip-1(gk466982)* nonsense alleles, both *jip-1* deletion alleles alone displayed a low penetrance of 2AWC^ON^ phenotype (Figure 1C, rows 3-6). In addition, *jip-1a/dE1-19 knock-out* re-capitulated the 2AWC^ON^ enhancement phenotype of *tir-1(ky388ts)* observed in *jip-1(vy6)* mutants (Figure 1C, rows 10-12). Together, these results suggest that the *vy6* mutation is a loss-of-function allele of *jip-1* in AWC asymmetry.

The transgene *jip-1p::jip-1a*, expressing *jip-1a* cDNA from a 3.3 kb *jip-1* promoter (Figure S2A), completely rescued the enhancement of 2AWC^ON^ penetrance from 99% to 41% in *tir-1(ky388ts); vy6* double mutants (Figure 1C, rows 10, 11, and 15). This result further supports that the 2AWC^ON^ enhancement phenotype of *tir-1(ky388ts); vy6* double mutants was caused by the identified missense mutation in *jip-1*.

We also tested the rescuing ability of different *jip-1* isoforms expressed from an AWC *odr-3* promoter. *odr-3p::jip-1a*, *odr-3p::GFP::jip-1a* (expressing GFP::JIP-1a fusion protein), and *odr-3p::jip-1d* (Figure S2A), rescued the 2AWC^ON^ enhancement phenotype of *tir-1(ky388ts); jip-1(vy6)* double mutants to a degree similar to *jip-1p::jip-1a* (Figure 1C, rows 10, 11, 15-18). However, *odr-3p::jip-1e* and *odr-3p::jip-1f* only slightly rescued the 2AWC^ON^ enhancement phenotype of *tir-1(ky388ts); jip-1(vy6)* double mutants (Figures 1C, rows 10, 11, 19, and 20; S2A). These results suggest that *jip-1a* and *jip-1d*, but not *jip-1e* or *jip-1f*, are essential for AWC asymmetry. These results are also consistent with the finding that the *vy6* mutation affects *jip-1a* and *jip-1d* but not *jip-1b/c/e/f* (Figures 1D and S2A). Furthermore, the *odr-3p::mJIP-1* transgene expressing mouse JIP-1 cDNA from the AWC *odr-3* promoter rescued the 2AWC^ON^ enhancement phenotype of *tir-1(ky388ts); vy6* double mutants (Figures 1C, rows 10, 11, and 21; S2A), similar to the rescuing ability of *odr-3p::jip-1a* (Figures 1C, rows 15 and 21) and *odr-3p::jip-1d* (Figures 1C, rows 18 and 21). This result suggests functional conservation between *C. elegans* and mammalian JIP-1 proteins.

In addition, we tested the rescuing ability of *odr-3p::GFP::jip-1a* for mislocalized TIR-1::TagRFP in *jip-1(vy6)* mutants. *odr-3p::GFP::jip-1a*, expressing GFP::JIP-1a in AWC, rescued the mislocalized TIR-1::TagRFP phenotype in *jip-1(vy6)* mutants (Figure 1B). Together, these results suggest that *jip-1a* mainly acts in AWC to regulate the synaptic TIR-1 localization in the AWC axon for promoting the AWC^OFF^ subtype.

### *jip-1a/d* is expressed in AWC neurons

To determine the expression pattern of *jip-1a* and *jip-1d* isoforms, the two isoforms essential for AWC asymmetry, we generated *mNG::SEC::jip-1a/d* and *mNG::jip-1a/d knock-in* animals by tagging the N-terminal end of endogenous JIP-1a/d protein isoforms with mNG::SEC or mNG using Cas9-triggered homologous recombination (DICKINSON *et al*. 2013; DICKINSON *et al*. 2015; DICKINSON AND GOLDSTEIN 2016) (Figure 1D). Since the self-excising cassette (SEC) contains transcriptional terminators, *mNG::SEC::jip-1a/d knock-in* animals are potentially loss-of-function mutants and transcriptional reporter strains of *jip-1a/d* isoforms. *mNG::jip-1a/d knock-in* animals, generated by SEC excision, are translational reporter strains of JIP-1a/d protein isoforms. Before the analysis of the expression pattern of *jip-1*, the localization pattern of TIR-1::TagRFP in AWC neurons and AWC asymmetry phenotypes were analyzed in *mNG::SEC::jip-1a/d knock-in* and *mNG::jip-1a/d knock-in* animals to determine the functionality of mNG::JIP-1 fusion proteins.

*mNG::SEC::jip-1a/d knock-in* animals had reduced TIR-1::TagRFP localization in the AWC axon and accumulation of TIR-1::TagRFP in the AWC cell body (Figures 1B), similar to the TIR-1::GFP or TIR-1::TagRFP localization phenotype caused by *jip-1(vy6)* and other *jip-1* loss-of-function mutant alleles (Figure 1A and 1B). These results suggest that *mNG::SEC::jip-1a/d knock-in* animals are loss-of-function mutants of *jip-1a/d* for synaptic localization of TIR-1 in the AWC axon. In contrast, *mNG::jip-1a/d knock-in* animals, with SEC excised, had a wild-type TIR-1::TagRFP localization pattern (Figure 1B), suggesting that mNG::JIP-1a/d fusion proteins are functional for the synaptic localization of TIR-1 in the AWC axon.

*mNG::SEC::jip-1a/d knock-in* animals displayed wild-type AWC asymmetry (Figure 1C, row 7). Similar to *jip-1(vy6)* and *jip-1a/dE1-19 knock-out* deletion allele, *mNG::SEC::jip-1a/d knock-in* greatly enhanced the 2AWC^ON^ phenotype of *tir-1(ky388ts)* from 22% to 97% at 15°C (Figures 1C, rows 9 and 13). However, *mNG::jip-1a/d knock-in* did not enhance the 2AWC^ON^ phenotype of *tir-1(ky388ts)* (Figure 1C, rows 9 and 14). Together, these results support that mNG::JIP-1a/d fusion proteins are functional in regulating the synaptic localization of TIR-1 in the AWC axon for AWC asymmetry.

*mNG::SEC::jip-1a/d knock-in* displayed mNG expression in numerous cells in the first-stage larvae’s (L1) head, body, and tail (Figure 2A). The broad expression pattern of *jip-1a/d* in the head is consistent with the expression of vertebrate *jip-1* in multiple brain regions, including the olfactory bulb (PELLET *et al*. 2000; SATO *et al*. 2011). mNG expression from *mNG::SEC::jip-1a/d knock-in* was also detected in AWC neurons (Figure 2A), consistent with the rescue data (Figure 1C, rows 10, 11, and 16), suggesting that *jip-1a* and *jip-1d* may act in AWC neurons to regulate the synaptic localization of TIR-1 for the specification of AWC^OFF^.

**Figure 2.**
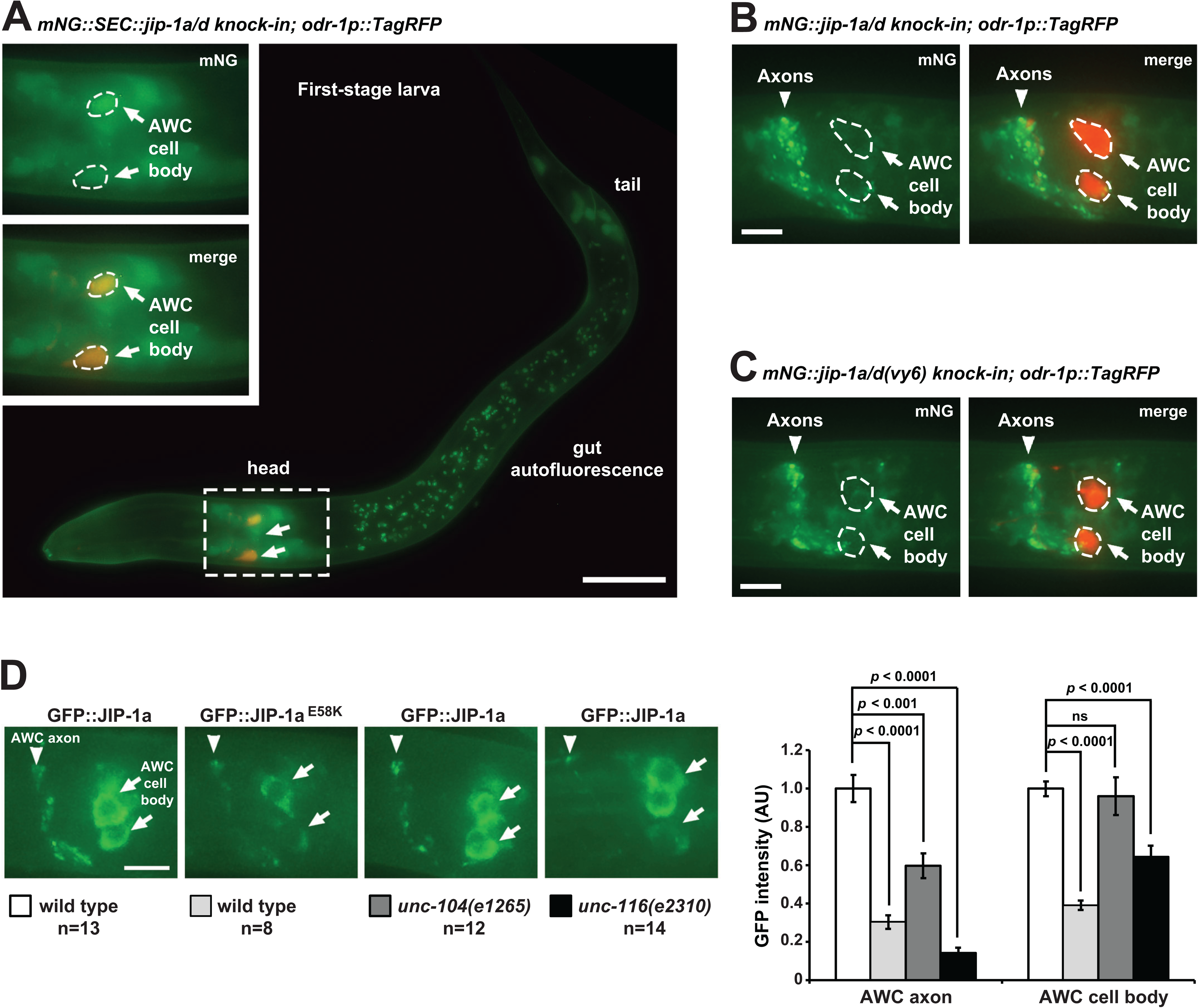
*jip-1a/d* is expressed in AWC neurons. (**A**) Representative images of *mNG::SEC::jip-1a/d knock-in* expression in a first-stage larva. The head area containing both AWC cells is outlined with dashed lines and magnified 2-fold in insets. AWC neurons were labeled with *odr-1p::TagRFP.* The AWC cell body is outlined with dashed lines. Green represents mNG; red labels TagRFP; yellow represents green signal overlapping with red. The anterior is left, and the ventral is down. Scale bar, 20 μm. (**B-C**) Representative images of *mNG::jip-1a/d knock-in* (B) and *mNG::jip-1a/d(vy6) knock-in* (C) expression in first-stage larvae. AWC neurons were labeled with *odr-1p::TagRFP.* The AWC cell body is outlined with dashed lines. The anterior is left, and the ventral is down. Scale bars, 5 μm. (**D**) Left panels: Images of GFP::JIP-1a and GFP::JIP-1a^E58K^, expressed from single copy insertion transgenes *odr-3p::GFP::jip-1a* and *odr-3p::GFP::jip-1a(vy6)*, respectively. Animals were imaged at the L1 stage. The anterior is left, and the ventral is down. Scale bars, 5 μm. Right panel: Quantification of GFP fluorescence intensity in the AWC axon and cell body at the L1 stage. AU, arbitrary unit. n, the total number of animals analyzed. Student’s *t*-test was used for statistical analysis. ns, not significant. Error bars, standard errors of the mean.

mNG::JIP-1a/d was localized in a punctate pattern in the axon processes of neurons in the head but was mostly excluded from the AWC cell body (Figure 2B). mNG::JIP-1a/d^E58K^, expressed from *mNG::jip-1a/d(vy6) knock-in*, displayed a reduced expression level in the axons of head neurons than mNG::JIP-1a/d (Figure 2B and 2C). To directly compare the subcellular localization pattern of JIP-1a and JIP-1a^E58K^ in AWC neurons, single-copy transgenes of *odr-3p::GFP::jip-1a*, which rescued *jip-1(vy6)* mutant phenotypes (Figure 1B and 1C, rows 10, 11, and 17), and *odr-3p::GFP::jip-1a(vy6)* were individually inserted on the same locus of a chromosome. Both GFP::JIP-1a, expressed from *odr-3p::GFP::jip-1a*, and GFP::JIP-1a^E58K^, expressed from *odr-3p::GFP::jip-1a(vy6)*, were localized in the AWC axon and cell body (Figure 2D). GFP::JIP-1a and GFP::JIP-1a^E58K^ displayed strong expression, in contrast to almost absent expression of mNG::JIP-1a/d knock-in and mNG::JIP-1a/d^E58K^ knock-in, in the AWC cell body, suggesting that the *odr-3* promoter used in the transgenes has a stronger activity than the endogenous *jip-1* promoter in AWC neurons (Figure 2B, 2C, and 2D). Consistent with the comparison between mNG::JIP-1a/d knock-in and mNG::JIP-1a/d^E58K^ knock-in, the fluorescence intensity of JIP-1a^E58K^ is significantly reduced in the AWC axon and cell body, compared to GFP::JIP-1a (Figure 2D). These results suggest that the *vy6* mutation may reduce the stability of the JIP-1a protein in AWC neurons.

### *jip-1a/d* is asymmetrically expressed at a higher level in the AWC^ON^ neuron

While mNG::SEC::JIP-1a/d was detected in both AWC neurons (Figures 2A and 3A), it was asymmetrically expressed at a higher level in the left AWC neuron (AWCL) or the right AWC neuron (AWCR) in a stochastic manner (Figure 3A and 3B). The stochastic asymmetry of *jip-1a/d* expression in AWC neurons is consistent with the random nature of AWC asymmetry. Asymmetric expression of *jip-1* in AWC was also observed in our initial assessment of *jip-1* expression pattern using *jip-1p::GFP*, in which GFP was expressed from the 3.3 kb *jip-1* promoter that rescued *jip-1(vy6)* mutant phenotype when driving the expression of *jip-1a* cDNA (Figures 1C, 3C, 3D, S2A). The percentage of animals with equivalent *jip-1p::GFP* expression level in AWCL and AWCR was significantly increased in *nsy-5(ky634)* (gap junction), *nsy-4(ky627)* (claudin), *unc-36(e251)* (regulatory subunit of calcium channels), and *unc-43(n1186)* (CaMKII) mutants (Figure 3D). UNC-2/EGL-19/UNC-36 calcium channels and UNC-43 CaMKII are required for the specification of AWC^OFF^ (TROEMEL *et al*. 1999; SAGASTI *et al*. 2001; BAUER HUANG *et al*. 2007), while NSY-5 gap junction protein and NSY-4 claudin-like protein act in parallel to inhibit the downstream calcium-regulated signaling pathway for the induction of the AWC^ON^ subtype (VANHOVEN *et al*. 2006; CHUANG *et al*. 2007). These results suggest that these AWC^ON^- and AWC^OFF^-promoting molecules, which act upstream of TIR-1, regulate asymmetric expression of *jip-1a/d* in AWC neurons.

**Figure 3.**
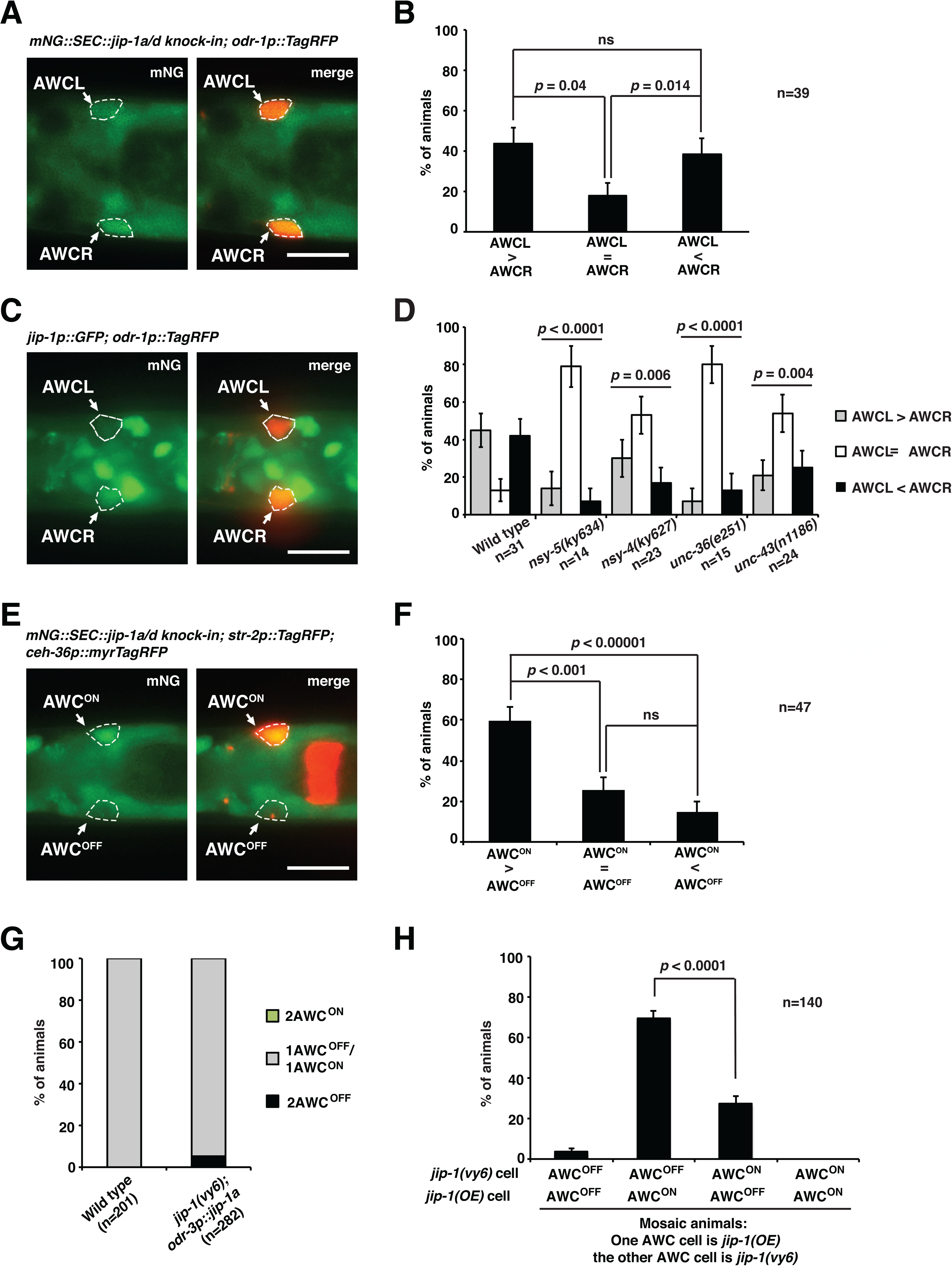
*jip-1a* is asymmetrically expressed at a higher level in AWC^ON^ and acts non-cell autonomously to promote AWC^OFF^. (**A, C**) Representative images of *mNG::jip-1a/d knock-in* (A) and *jip-1p::GFP* (C) expression at a higher level in the AWCR neuron than in AWCL at the L1 stage (ventral view). Both AWCL and AWCR were marked by *odr-1p::TagRFP.* (**B**, **D**) Quantification of asymmetric *mNG::jip-1a/d knock-in* (B) and *jip-1p::GFP* (D) expression in AWCL and AWCR neurons at the L1 stage. No significant difference was observed between AWCL>AWCR and AWCL<AWCR in wild-type animals. (**E**) Images of *mNG::jip-1a/d knock-in* expression at a higher level in AWC^ON^ than in AWC^OFF^ in an L1 animal (dorsal view). AWC^ON^ was marked by *str-2p::TagRFP* and *ceh-36p::myrTagRFP*, while AWC^OFF^ was only marked by *ceh-36p::myrTagRFP*. (**F**) Quantification of asymmetric expression of *mNG::jip-1a/d knock-in* in AWC^ON^ and AWC^OFF^ at the L1 stage. (**G**) Quantification of AWC asymmetry phenotypes in wild type and *jip-1(vy6)* mutants containing the extrachromosomal transgene *odr-3p::jip-1a(OE); odr-1p::DsRed* in adults. (**H**) Quantification of AWC phenotypes in *jip-1(vy6)* mosaic animals containing the extrachromosomal transgene *odr-3p::jip-1a(OE)* in only one AWC neuron, inferred by the presence of the co-injected *odr-1p::DsRed* AWC marker. The data was obtained from a subset of animals scored in (G). The AWC cell body is outlined with dashed lines (A, C, E). Scale bars, 10 μm. *P*-values were calculated using a *Z*-test (B, F, H) and Fisher’s exact test (D). ns, not significant. Error bars represent the standard error of the proportion. n, total number of animals scored.

In addition, mNG::SEC::JIP-1a/d was expressed at a significantly higher expression level in the AWC^ON^ neuron than the AWC^OFF^ neuron in most animals (Figure 3E and 3F). These results suggest the hypothesis that *jip-1a/d* mainly acts in AWC^ON^ to non-cell autonomously promote the AWC^OFF^ subtype.

### *jip-1* acts non-cell autonomously in AWC^ON^ to promote AWC^OFF^

Expression of *jip-a* in AWC from the *odr-3* promoter significantly rescued the 2AWC^ON^ enhancement phenotype of *tir-1(ky388ts); jip-1(vy6)* double mutants (Figure 1C, rows 10, 11, and 16) and the mislocalized TIR-1 phenotype in *jip-1(vy6)* mutants (Figure 1B), suggesting that *jip-1a* mainly acts in AWC cells to regulate the synaptic TIR-1 localization for promoting the AWC^OFF^ subtype. Mosaic animals in which *jip-1a* activity is different between the two AWC neurons were used to determine whether *jip-1a* acts in AWC^ON^, AWC^OFF^, or both to promote the AWC^OFF^ subtype. Expression of *odr-3p::jip-1a* extrachromosomal transgenes in both AWC cells caused a slight 2AWC^OFF^ phenotype in *vy6* mutants (Figure 3G). Spontaneous loss of the extrachromosomal array resulted in mosaic animals in which only one of the AWC neurons retained the *odr-3p*::*jip-1a* transgene (inferred by the co-injected AWC marker *odr-1p::DsRed*), and the other AWC cell remained *jip-1(vy6)*. In most mosaic animals, the AWC cell expressing *odr-3p::jip-1a* became AWC^ON^, and the *jip-1(vy6)* AWC cell became AWC^OFF^ (Figure 3H). These results suggest that *jip-1a* acts largely in AWC^ON^ to non-cell autonomously promote AWC^OFF^, similar to the previously identified non-cell-autonomous role of UNC-104/kinesin-3 (KIF1A) and UNC-116/kinesin-1 motor proteins in the specification of AWC^OFF^ (CHANG *et al*. 2011; KHALID *et al*. 2026).

### JIP-1 regulates the dynamic trafficking of TIR-1 in the AWC axon

We previously showed that TIR-1::GFP, expressed from the transgene *odr-3p::tir-1::GFP*, is dynamically transported in the AWC axon in a manner dependent on UNC-104/kinesin-3 (KIF1A) and UNC-116/kinesin-1 motor proteins (CHANG *et al*. 2011; KHALID *et al*. 2026). To examine the role of JIP-1 in the dynamic trafficking of TIR-1 in the AWC axon, the *odr-3p::tir-1::GFP* transgene was used for time-lapse imaging of TIR-1::GFP in wild type and *jip-1(vy6)* mutants (Figure 4A). In wild type, TIR-1::GFP moved in both anterograde (from cell body to axon) and retrograde (from axon to cell body) directions with almost equal ratios (Figure 4B). In *jip-1(vy6)* mutants, similar to *unc-104(e1265)* and *unc-116(e2310)* mutants, the ratio of anterograde movement of TIR-1::GFP was significantly reduced (CHANG *et al*. 2011; KHALID *et al*. 2026) (Figure 4B). The average velocity of TIR-1::GFP between anterograde and retrograde movement directions was not significantly different in wild type or *jip-1(vy6)* mutants (Figure 4C). However, the average velocity of the TIR-1::GFP movement in both anterograde and retrograde directions was significantly increased in *jip-1(vy6)* mutants, compared to the wild type (Figure 4C). Together, these results are consistent with reduced localization of TIR-1 in the AWC axon and accumulation of TIR-1 in the AWC cell body in *jip-1(vy6)* mutants. These results also suggest that JIP-1 promotes anterograde transport of TIR-1 in the AWC axon by regulating the movement’s directionality and relative velocity.

**Figure 4.**
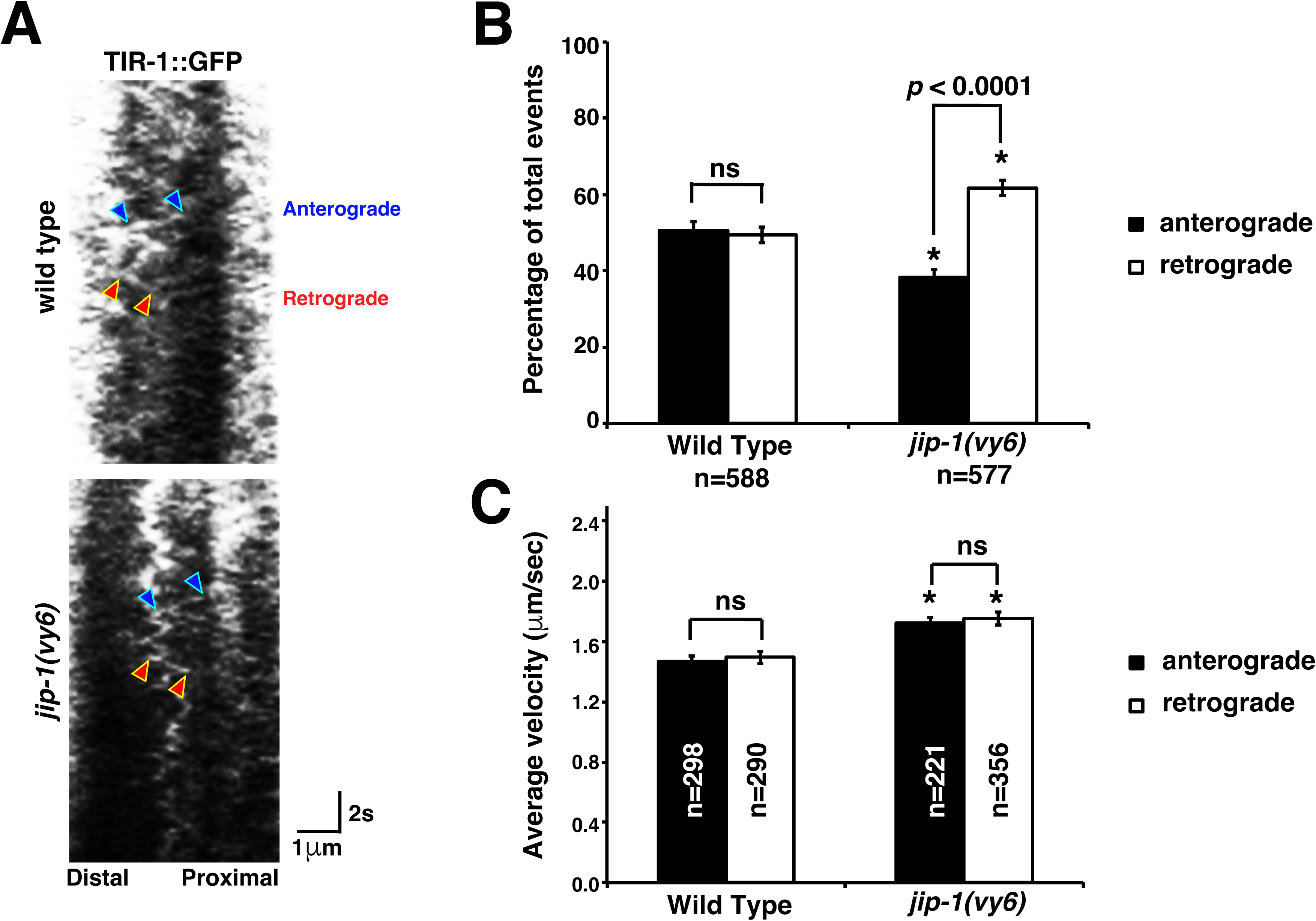
JIP-1 regulates the dynamic trafficking of TIR-1 in the AWC axon. (A) Representative kymographs of TIR-1::GFP movement in the AWC axon of wild type and *jip-1(vy6)* mutants at the L1 stage. Blue arrowheads indicate anterograde movement (from cell body to axon), and red arrowheads indicate retrograde movement (from axon to cell body). (**B, C**) Quantification of the percentage of anterograde and retrograde TIR-1::GFP trafficking events (B) and the average velocity of trafficking events (C). Asterisks indicate significant differences (*p* < 0.0001) in the same direction of movement between wild type and *jip-1(vy6)* mutants. *P*-values were determined by a *Z*-test (B) or Student’s *t*-test (C). Error bars represent the standard error of the proportion (B) or the standard error of the mean (C). ns, not significant. n, number of trafficking events.

### UNC-104 and UNC-116 are required for the dynamic trafficking of JIP-1 in the AWC axon

We also examined whether JIP-1 is dynamically transported in the AWC axon using time-lapse imaging of GFP::JIP-1a and GFP::JIP-1a^E58K^, expressed from transgenes driven by the AWC *odr-3* promoter (Figure S3A). In the wild type, GFP::JIP-1a, like TIR-1::GFP, is dynamically transported in the AWC axon in both anterograde and retrograde directions with equivalent ratios of total events (Figure S3B). GFP::JIP-1a^E58K^, compared to GFP::JIP-1a, had a significantly decreased ratio of anterograde movement (Figure S3B). The average velocity of GFP::JIP-1a and GFP::JIP-1a^E58K^ between anterograde and retrograde movement directions was not significantly different (Figure S3C). These results are consistent with the decreased localization of GFP::JIP-1a^E58K^, compared to GFP::JIP-1a, in the AWC axon (Figure 2D).

GFP::JIP-1a displayed a significantly lower ratio of anterograde movement than retrograde movement in *unc-104(e1265)* and *unc-116(e2310)* mutants, compared to wild type (CHANG *et al*. 2011; KHALID *et al*. 2026) (Figure S3B). These results are consistent with the decreased localization of GFP::JIP-1a in the AWC in *unc-104(e1265)* and *unc-116(e2310)* mutants (Figure 2D). These results also suggest that UNC-104/kinesin-3 (KIF1A) and UNC-116/kinesin-1 motor proteins promote the anterograde transport of JIP-1 in the AWC axon.

### JIP-1 is localized in close proximity to UNC-104/kinesin-3 and UNC-116/kinesin-1 in the AWC axon

Our genetic mosaic analyses suggest that *jip-1*, like *unc-104* and *unc-116,* acts largely in AWC^ON^ to non-cell autonomously promote AWC^OFF^ (CHANG *et al*. 2011; KHALID *et al*. 2026) (Figure 3G and 3H). Time-lapse imaging also reveals a role of JIP*-*1, like UNC*-*104 and UNC-116, in the dynamic trafficking of TIR-1, and a role of UNC*-*104 and UNC-116 in the dynamic transport of JIP-1 in the AWC axon (CHANG *et al*. 2011; KHALID *et al*. 2026) (Figures 4 and S3). In addition, our previous study showed that UNC-104 and UNC-116 displayed almost completely overlapping expression patterns along the AWC axons (CHANG *et al*. 2011; KHALID *et al*. 2026). Together, these results suggest that JIP-1 may be located close to UNC-104 and UNC-116 in the AWC axon to non-cell autonomously promote anterograde transport of TIR-1. To test this, we examined the localization patterns of these proteins expressed from fluorescently tagged transgenes driven by the AWC *odr-3* promoter in the AWC axon.

GFP::JIP-1a and UNC-104::TagRFP displayed overlapping expression patterns along the AWC axon (Figure 5A). Similarly, TagRFP::JIP-1a and UNC-116::GFP overlapped along the AWC axon (Figure 5B). In addition, JIP-1, like UNC-104 and UNC-116 (CHANG *et al*. 2011; KHALID *et al*. 2026), was localized adjacent to TIR-1 along the AWC axon (Figure 5C). These results suggest that JIP-1, UNC-104/kinesin-3 (KIF1A), and UNC-116/kinesin-1 proteins may function in proximity to regulate the dynamic transport of TIR-1 in the AWC axon for promoting AWC^OFF^.

**Figure 5.**
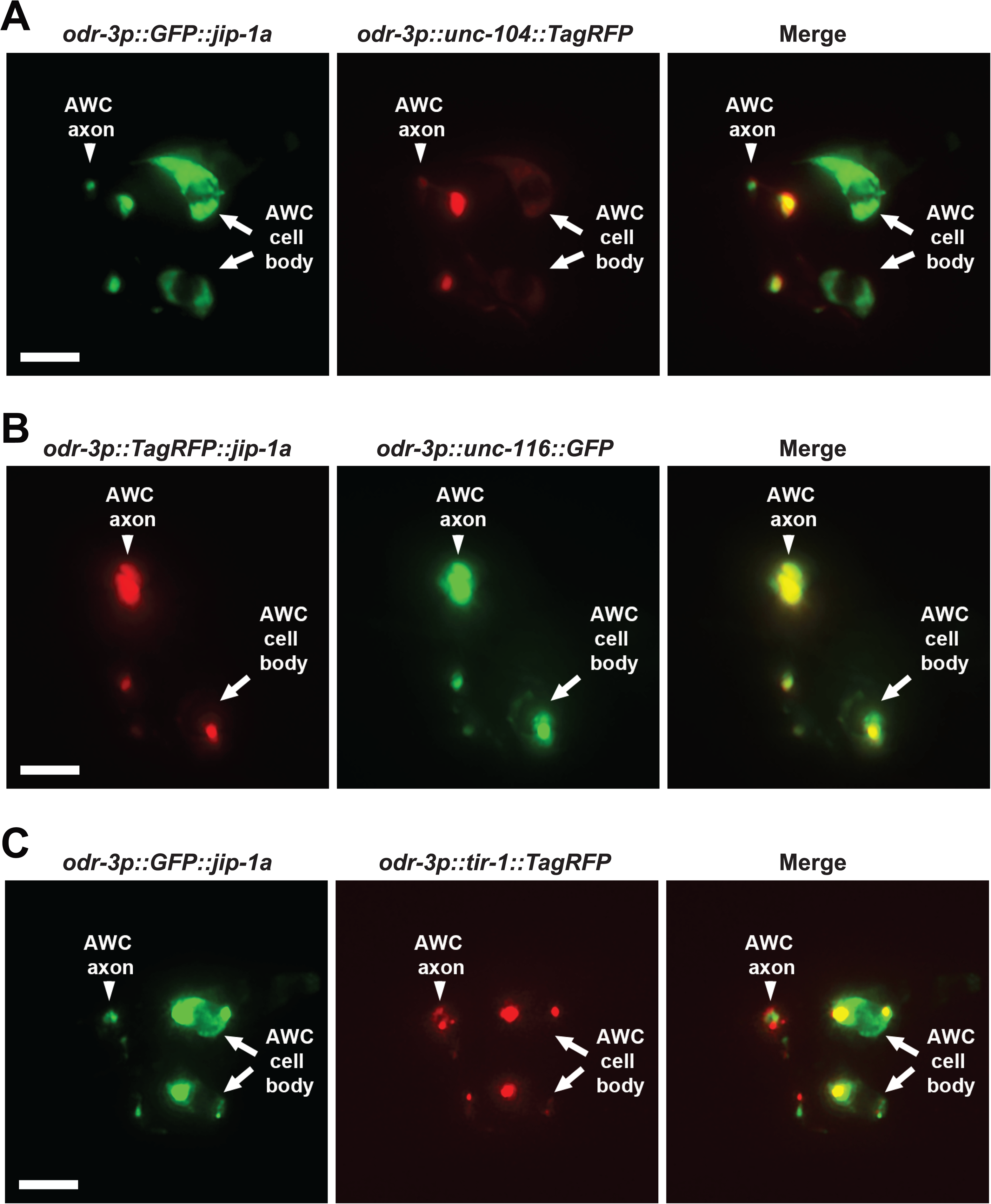
JIP-1 colocalized with UNC-104/kinesin-3 and UNC-116/kinesin-1 and is localized adjacent to TIR-1 in the AWC axons. **(A)** Images of a wild-type L1 animal expressing *odr-3p::GFP::jip-1a* and *odr-3p::unc-104::TagRFP* in AWC neurons. **(B)** Images of a wild-type L1 animal expressing *odr-3p::TagRFP::jip-1a* and *odr-3p::unc-116::GFP* in AWC neurons. (**C**) Images of a wild-type L1 animal expressing *odr-3p::GFP::jip-1a* and *odr-3p::tir-1::TagRFP* in AWC neurons. Scale bar, 5μm. Anterior to the left and ventral at the bottom.

## Discussion

Here, we identify the role of the JNK-interacting protein JIP-1 in the stochastic choice of asymmetric olfactory neuron subtypes in *C. elegans* from an unbiased forward genetic screen. JIP-1 mediates synaptic localization of the calcium-signaling scaffold protein TIR-1/SARM1 in the AWC axon to promote the AWC^OFF^ subtype. This study implicates JIP-1 in establishing left-right patterning and in stochastic cell-identity choice. As *C. elegans* JIP-1 is functionally conserved with mammalian JIP-1 proteins, this process may be conserved in establishing left-right asymmetry and stochastic cell identity in mammals.

We previously proposed that the UNC-104/kinesin-3 and UNC-116/kinesin-1 may work cooperatively to transport some unknown presynaptic factor(s) (illustrated as Y in the Figure 6 model) in the future AWC^ON^ cell that trans-synaptically regulates the dynamic trafficking of the TIR-1/SARM1 signaling complex to postsynaptic regions of the contralateral AWC axons in promoting the AWC^OFF^ subtype (CHANG *et al*. 2011; HSIEH *et al*. 2014; KHALID *et al*. 2026).

**Figure 6.**
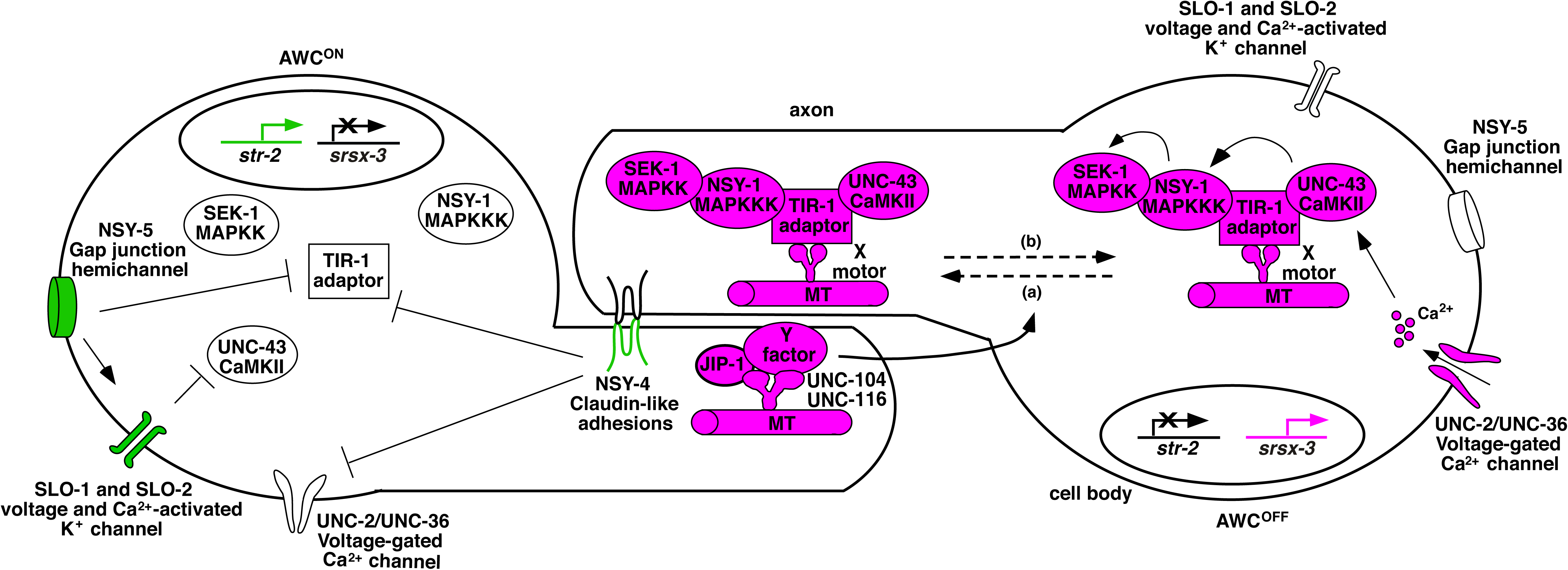
Model of JIP-1 function in AWC subtype specification. The specification of the two AWC subtypes is stochastic. In this model, AWC^ON^ is illustrated on the left, and AWC^OFF^ is on the right. Molecules in magenta promote AWC^OFF^, green promotes AWC^ON^, and white indicates less active or inactive molecules. In the default AWC^OFF^ subtype, calcium entry through UNC-2/UNC-36 voltage-gated calcium channels activates UNC-43/CaMKII, leading to the assembly of a calcium-signaling complex consisting of UNC-43/CaMKII, the TIR-1/SARM1 adaptor protein, and NSY-1/MAPKKK. The assembly of the calcium-signaling complex allows UNC-43/CaMKII to phosphorylate NSY-1/MAPKKK and then NSY-1/MAPKKK to phosphorylate SEK-1/MAPKK. Microtubules (MT) mediate the anterograde (a) and retrograde (b) transport of the TIR-1 signaling complex in the AWC axon via unidentified motor proteins (illustrated as X here). UNC-104 and UNC-116 kinesin motor proteins work together with JIP-1 in the contralateral AWC^ON^ cell to transport an unknown Y molecule, which non-cell autonomously regulates the dynamic transport of the TIR-1 signaling complex in the AWC^OFF^ cell to specify the AWC^OFF^ subtype. In the induced AWC^ON^ subtype, NSY-4 claudin-like adhesions act in parallel with NSY-5 gap junctions and SLO BK potassium channels to inhibit calcium channel-mediated signaling, leading to de-repression of *str-2* expression.

This study reveals a role for JIP-1, like UNC-104 and UNC-116, in promoting synaptic transport of TIR-1, which is required for the AWC^OFF^ subtype. Like *unc-104* and *unc-116*, *jip-1* acts in AWC^ON^ to non-cell autonomously promote AWC^OFF^. Our results support a model in which JIP-1 cooperates with UNC-104 and UNC-116 to non-cell autonomously regulate the dynamic trafficking of TIR-1 along the AWC axon for AWC^OFF^ specification. Our study extends the previous model of AWC asymmetry by identifying JIP-1 as a potential adaptor protein linking the kinesin proteins UNC-104 and UNC-116 to the Y factor in AWC^ON^, which non-cell autonomously regulates synaptic trafficking of the TIR-1 signaling complex to specify the AWC^OFF^ subtype (Figure 6).

The proposed role of JIP-1 in asymmetric AWC neuronal subtype choice is consistent with previously identified roles of JIPs as adaptors linking molecular motors to cargos in axonal transport. The phosphorylation of JIP-1 acts as a molecular switch between its association with anterograde and retrograde motile complexes, thereby regulating the directionality of the axonal transport of amyloid precursor protein in cultured mouse primary dorsal root ganglion sensory neurons (FU AND HOLZBAUR 2013). The *Drosophila* JIP-1 protein APLIP1 is associated with kinesin-1-driven anterograde and dynein-driven retrograde vesicle transport (HORIUCHI *et al*. 2005). The *Drosophila* JIP3 protein Sunday Driver mediates the axonal transport of vesicles by directly interacting with kinesin-1 (BOWMAN *et al*. 2000). The *C. elegans* JIP-3 protein UNC-16 functions as an adaptor to link kinesin-1 with dynein for kinesin-1-dependent anterograde transport of dynein and, in turn, promotes dynein-mediated retrograde transport of cargos in touch and motor neurons (ARIMOTO *et al*. 2011). *C. elegans* UNC-16/JIP-3 also interacts with kinesin-1 to regulate the transport or localization of synaptic vesicle components in motor neurons (BYRD *et al*. 2001; SAKAMOTO *et al*. 2005). In cultured rat hippocampal neurons, JIP-3 links kinesin-1 with TrkB receptors to drive anterograde transport of TrkB into distal axons and consequently facilitates BDNF (brain-derived neurotrophic factor)-induced retrograde signaling (HUANG *et al*. 2011).

Similar to mammalian *jip* genes, which generate multiple transcript variants through alternative splicing (KIM *et al*. 1999), the two *C. elegans jip* genes, *jip-1* and *jip-3* (*unc-16*), have 6 and 16 alternatively spliced isoforms, respectively (wormbase.org). All *jip* variants identified in mice and rats are preferentially expressed in the brain (KIM *et al*. 1999). The biological significance of multiple *jip* transcript variants has been poorly elucidated. Our results show that *jip-1a* and *jip-1d*, but not *jip-1b/c/e/f* (Hsieh, Y.-W., and Chuang, C.-F., unpublished), are expressed in AWC cells and play an essential role in the synaptic localization of TIR-1 for the specification of AWC^OFF^. Our study reveals cell-specific expressions and functions of *jip-1* isoforms.

JIPs have been implicated in a variety of neuronal development and functions. JIP-1 regulates axonal growth of mouse cortical neurons (DAJAS-BAILADOR *et al*. 2008; DAJAS-BAILADOR *et al*. 2014), axon guidance of mouse telencephalic commissures (HA *et al*. 2005), and axonal regeneration of mouse primary dorsal root ganglion neurons (BARNAT *et al*. 2010). JIP-3 is required for axon elongation of rat hippocampal and cortical neurons (SUN *et al*. 2013), axon branching of rat cerebellar granule neurons (BILIMORIA *et al*. 2010), and synaptic membrane trafficking of *C. elegans* motor neurons (BROWN *et al*. 2009). Our findings reveal a role for JIP-1 in the kinesin-dependent trafficking of cell-specific Ca^2+^ signaling proteins to synaptic regions, thereby enabling stochastic choice of olfactory neuron subtypes.

## Materials and Methods

### Strains and transgenes

The wild-type *C. elegans* strain is N2, Bristol variety. Strains were cultured by standard methods (BRENNER 1974). A list of strains and transgenes is included in Supplemental Materials and Methods.

### Isolation of *jip-1(vy6)* mutants

A forward genetic screen was performed as previously described (BRENNER 1974). Integrated *odr-3p::tir-1::GFP* transgenic strain P_0_ was treated with EMS, five F_1_ progenies were picked onto single plates, and F_2_ were screened for defective TIR-1::GFP localization in the AWC axon using a Zeiss compound fluorescence microscope. The *vy6* mutation was identified from a screen of 2,000 genomes.

### Whole genome sequencing

The one-step whole-genome sequencing and SNP mapping strategy (DOITSIDOU *et al*. 2010) was used to identify the *vy6* mutation with an Illumina GA2x sequencing platform and 100-nucleotide reads. CloudMap software was used to analyze sequencing results as previously described (MINEVICH *et al*. 2012).

### Supplemental Information Appendix

Supplemental information includes supplemental materials and methods, supplemental references, supplemental figure legends, and supplemental figures S1-S3.

## Data availability

All data discussed in the paper will be available to readers.

## Acknowledgments

We thank Alex Boyanov, Oliver Hobert, Dan Dickinson, Bob Goldstein, WormBase, and the *C. elegans* Genetic Center (funded by the NIH Office of Research Infrastructure Programs P40 OD010440) for assistance, strains, reagents, protocols, and/or databases. We also thank Eleana Liu for comments on the manuscript. This work was supported by a Whitehall Foundation Research Award (C.-F.C.), Alfred P. Sloan Research Fellowship (C.-F.C.), and the National Institutes of Health (5R01GM098026-05 to C.-F.C.).

## Author Contributions

Conceptualization, Y.-W.H. and C.-F.C.; Methodology, Y.-W.H., C.S., S.Y., J.Y., and C.-F.C.; Investigation, Y.-W.H., C.S., S.Y., J.Y., and C.-F.C.; Formal Analysis, Y.-W.H., C.S., S.Y., J.Y., and C.-F.C.; Writing, Y.-W.H. and C.-F.C.; Funding Acquisition, C.-F.C.; Supervision, C.-F.C.

## Declaration of Interests

The authors declare no competing interests.

## Supplemental Materials and Methods

### Strains and transgenes

Animal protocols approved by the Office of Animal Care and Institutional Biosafety Committees at the University of Illinois Chicago were followed. Hermaphrodites of *C. elegans* were analyzed and imaged.

#### Mutants

*nsy-5(ky634)* I (CHUANG *et al*. 2007)

*jip-1(vy6)* II (This study)

*jip-1(gk133506)* II (THOMPSON *et al*. 2013)

*jip-1(gk466982)* II (THOMPSON *et al*. 2013)

*ttTi5605* II (Frokjaer-Jensen *et al*. 2008; Frokjaer-Jensen *et al*. 2012)

*unc-104(e1265)* II (KUMAR *et al*. 2010)

*tir-1(ky388ts)* III (Chuang and Bargmann 2005)

*unc-36(e251)* III (BRENNER 1974)

*unc-116(e2310)* III (PATEL *et al*. 1993)

*unc-119(ed3)* III (Frokjaer-Jensen *et al*. 2008; Frokjaer-Jensen *et al*. 2012)

*nsy-4(ky627)* IV (VANHOVEN *et al*. 2006)

*unc-43(n1186lf)* IV (Park and Horvitz 1986)

*cxTi10816* IV (Frokjaer-Jensen *et al*. 2008; Frokjaer-Jensen *et al*. 2012)

#### mNG knock-in strains

*jip-1(vy260 [mNG::SEC::jip-1a/d knock-in])* II (this study)

*jip-1(vy260 vy263 [mNG::jip-1a/d knock-in])* II (this study)

*jip-1(vy286 vy290 [mNG::jip-1a/d(vy6) knock-in])* II (this study)

#### mNG knock-out strains

*jip-1(vy308 [jip-1a/dE1-19 knock-out])* II (this study)

*jip-1(vy309 [jip-1a/dE12-19 knock-out])* II (this study)

#### Integrated transgenes

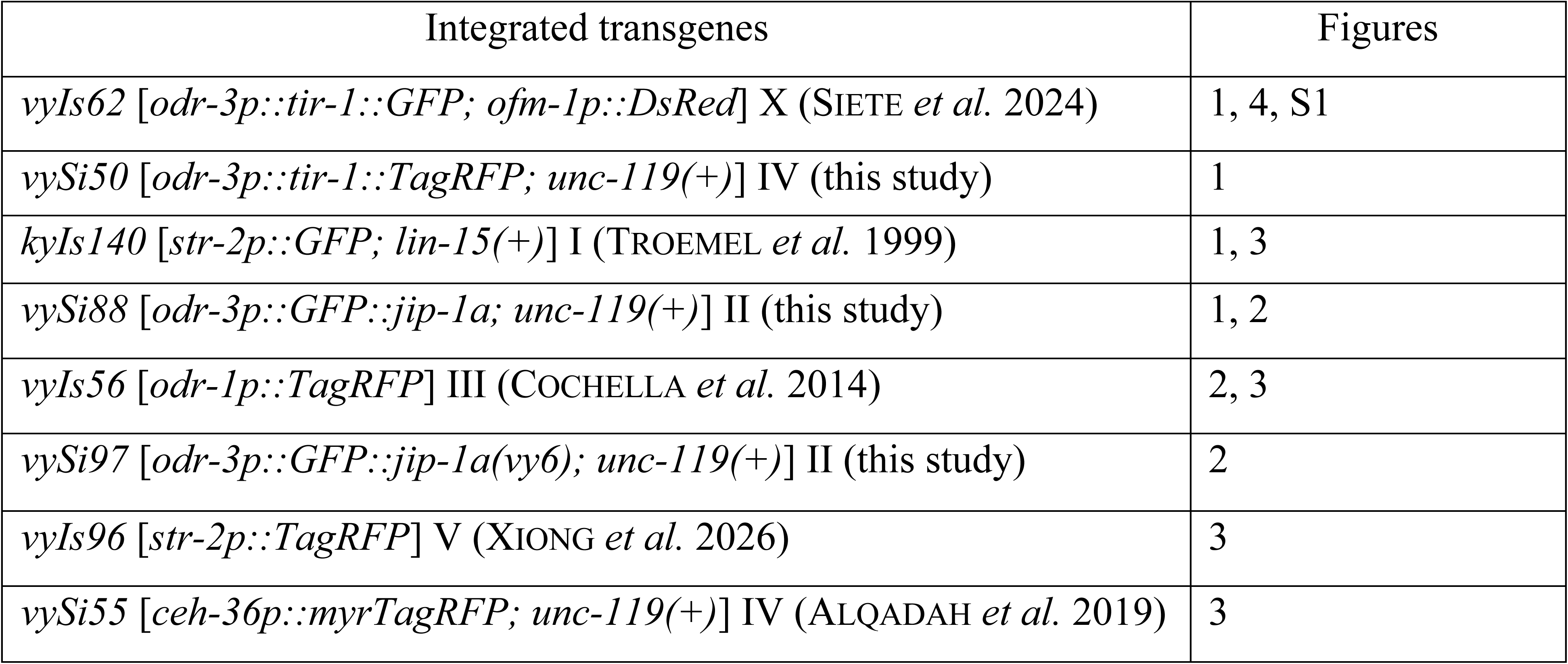

#### Extrachromosomal arrays

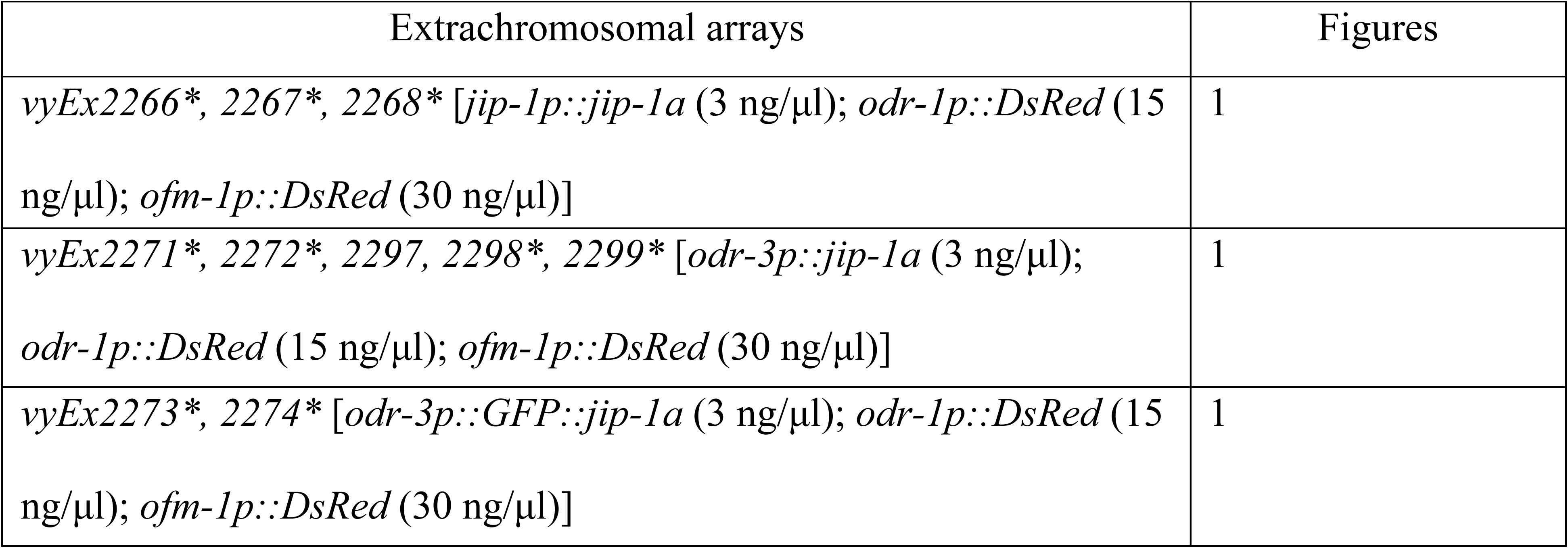
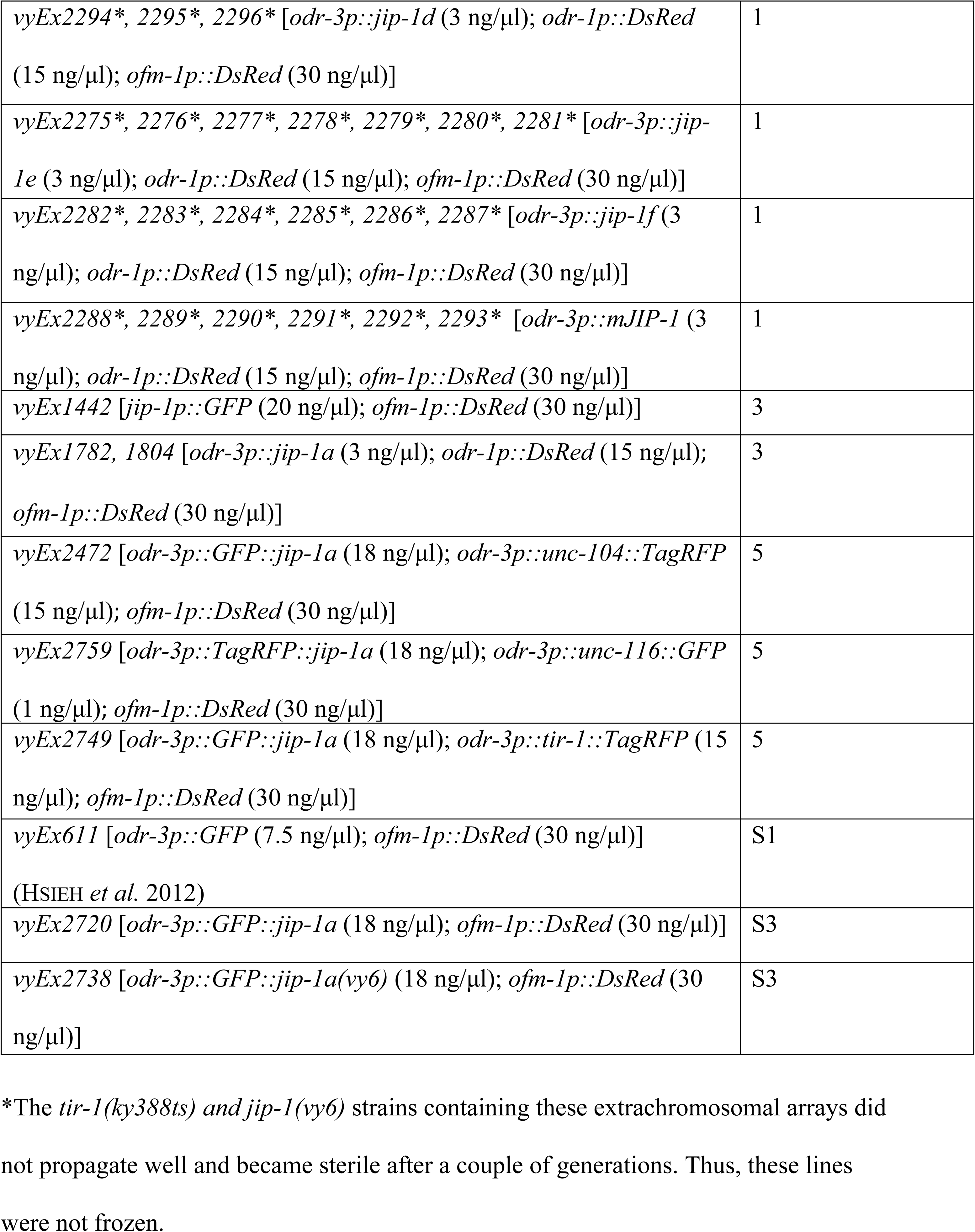

| Extrachromosomal arrays | Figures |
| --- | --- |
| *vyEx2266\*, 2267\*, 2268\* [jip-1p::*jip-1a* (3 ng/μl); odr-1p::*DsRed* (15 ng/μl); ofm-1p::*DsRed* (30 ng/μl)]* | 1 |
| *vyEx2271\*, 2272\*, 2297, 2298\*, 2299\* [odr-3p::*jip-1a* (3 ng/μl); odr-1p::*DsRed* (15 ng/μl); ofm-1p::*DsRed* (30 ng/μl)]* | 1 |
| *vyEx2273\*, 2274\* [odr-3p::*GFP::jip-1a* (3 ng/μl); odr-1p::*DsRed* (15 ng/μl); ofm-1p::*DsRed* (30 ng/μl)]* | 1 |
| vyEx2294*, 2295*, 2296* [*odr-3p::jip-1d* (3 ng/μl); *odr-1p::DsRed* (15 ng/μl); *ofm-1p::DsRed* (30 ng/μl)] | 1 |
| vyEx2275*, 2276*, 2277*, 2278*, 2279*, 2280*, 2281* [*odr-3p::jip-1e* (3 ng/μl); *odr-1p::DsRed* (15 ng/μl); *ofm-1p::DsRed* (30 ng/μl)] | 1 |
| vyEx2282*, 2283*, 2284*, 2285*, 2286*, 2287* [*odr-3p::jip-1f* (3 ng/μl); *odr-1p::DsRed* (15 ng/μl); *ofm-1p::DsRed* (30 ng/μl)] | 1 |
| vyEx2288*, 2289*, 2290*, 2291*, 2292*, 2293* [*odr-3p::mJIP-1* (3 ng/μl); *odr-1p::DsRed* (15 ng/μl); *ofm-1p::DsRed* (30 ng/μl)] | 1 |
| vyEx1442 [*jip-1p::GFP* (20 ng/μl); *ofm-1p::DsRed* (30 ng/μl)] | 3 |
| vyEx1782, 1804 [*odr-3p::jip-1a* (3 ng/μl); *odr-1p::DsRed* (15 ng/μl); *ofm-1p::DsRed* (30 ng/μl)] | 3 |
| vyEx2472 [*odr-3p::GFP::jip-1a* (18 ng/μl); *odr-3p::unc-104::TagRFP* (15 ng/μl); *ofm-1p::DsRed* (30 ng/μl)] | 5 |
| vyEx2759 [*odr-3p::TagRFP::jip-1a* (18 ng/μl); *odr-3p::unc-116::GFP* (1 ng/μl); *ofm-1p::DsRed* (30 ng/μl)] | 5 |
| vyEx2749 [*odr-3p::GFP::jip-1a* (18 ng/μl); *odr-3p::tir-1::TagRFP* (15 ng/μl); *ofm-1p::DsRed* (30 ng/μl)] | 5 |
| vyEx611 [*odr-3p::GFP* (7.5 ng/μl); *ofm-1p::DsRed* (30 ng/μl)]<br>(HSIEH *et al.* 2012) | S1 |
| vyEx2720 [*odr-3p::GFP::jip-1a* (18 ng/μl); *ofm-1p::DsRed* (30 ng/μl)] | S3 |
| vyEx2738 [*odr-3p::GFP::jip-1a(vy6)* (18 ng/μl); *ofm-1p::DsRed* (30 ng/μl)] | S3 |

\*The *tir-1(ky388ts)* and *jip-1(vy6)* strains containing these extrachromosomal arrays did not propagate well and became sterile after a couple of generations. Thus, these lines were not frozen.

### Plasmid construction

*pCFJ356::odr-3p::tir-1::TagRFP* was made by subcloning 7198 bp of *odr-3p::tir-1::TagRFP* into the pCFJ356 vector (FROKJAER-JENSEN *et al*. 2012).

*jip-1p::jip-1a* was generated by subcloning 3003 bp of *jip-1a* cDNA into a vector containing 3298 bp of the *jip-1* promoter.

*odr-3p::jip-1a* was generated by subcloning 3003 bp of *jip-1a* cDNA into a vector containing the *odr-3* promoter.

*odr-3p::jip-1d* was generated by deleting the 15 bp sequence (ccttcttctctccag) specific for *jip-1a/e/f* isoforms from *odr-3p::jip-1a* using Q5 site-directed mutagenesis kit (New England Biolabs)

*odr-3p::jip-1e* was generated by subcloning 1581 bp of *jip-1e* cDNA into a vector containing the *odr-3* promoter using Gibson assembly (New England Biolabs).

*odr-3p::jip-1f* was generated by subcloning 1119 bp of *jip-1f* cDNA into a vector containing the *odr-3* promoter using Gibson assembly (New England Biolabs).

*odr-3p::GFP::jip-1a* was generated by subcloning GFP into the 5’ end of *jip-1a* cDNA in the *odr-3*p*::jip-1a* vector using Gibson assembly (New England Biolabs).

*odr-3p::GFP::jip-1a(vy6)* was generated by subcloning GFP into the 5’ end of *jip-1a(vy6)* in the *odr-3*p*::jip-1a(vy6*) vector using Gibson assembly (New England Biolabs).

*odr-3p::mJIP-1* was generated by subcloning 2094 bp of mJIP-1(MAPK8IP1) cDNA into a vector containing the *odr-3* promoter.

*pAB1.1*::*odr-3p::GFP::jip-1a* was made by subcloning 7414 bp of *odr-3p::GFP::jip-1a* into the *pAB1.1* vector (ALQADAH *et al*. 2016).

*pAB1.1*::*odr-3p::GFP::jip-1a(vy6)* was made by subcloning 7414 bp of *odr-3p::GFP::jip-1a(vy6)* was subcloned into the *pAB1.1* vector (ALQADAH *et al*. 2016).

*odr-3p::TagRFP::jip-1a* was generated by subcloning 1643 bp of *TagRFP* into a vector containing the *odr-3p* promoter and *jip-1a* cDNA.

sgRNA constructs for Cas9-triggered homologous recombination were made by subcloning the sgRNA fragment into the *pDD162 eft-3p::Cas9::empty sgRNA* vector as previously described (DICKINSON *et al*. 2013; DICKINSON *et al*. 2015; DICKINSON AND GOLDSTEIN 2016) (Addgene #47549 from Bob Goldstein’s lab) using the Q5 site-directed mutagenesis kit (New England Biolabs).

Homology repair template constructs for Cas9-triggered homologous recombination were generated by subcloning 5’ and 3’ homology regions, generated by PCR, into the pDD268 mNG::SEC::3xFLAG vector (DICKINSON *et al*. 2013; DICKINSON *et al*. 2015; DICKINSON AND GOLDSTEIN 2016) using Gibson Assembly (New England Biolabs). One or both guanine nucleotides of the PAM (protospacer adjacent motif, NGG motif) site, if present in the homology fragment of the repair template construct, were mutated using a Q5 site-directed mutagenesis kit (New England Biolabs).

The sequence of the gRNA fragment cloned into *pDD162 eft-3p::Cas9::empty sgRNA* vector, primers used to PCR amplify homology regions, and the location of the mutated nucleotide(s) of the PAM site are listed below for individual knock-in and knock-out strains.

#### mNG::SEC::jip-1a/d knock-in

*eft-3p::Cas9::U6p::jip-1p sgRNA2* (GCCGACGGTTATTCGGAGGGT)

1759 bp of 5’ homology region

Forward primer: GTGAAATGAGGCGCTGAACTTG

Reverse primer: CGATTTGTGCTGAAATTTTTAAGACTTAAAATTTCAAG

1558 bp of 3’ homology region

Forward primer: ATGAGCGCATTTGAATGTCGAAAATG

Reverse primer: CTGGCAAATATTGGTCTGACGGACTAC

The GG nucleotides of the PAM site, located at 100 bp and 101 bp downstream of the *jip-1a/d* start codon within the 3’ homology region, were mutated to CC to generate a silent mutation.

#### mNG::SEC::jip-1a/d (vy6) knock-in

repair template construct was generated by mutagenizing *mNG::SEC::jip-1a/d knock-in* repair template construct using Q5 site-directed mutagenesis kit (New England Biolabs) with forward primer (AATTTCTGGTTCATCCGTGTCTTCG) and reverse primer (TGGCGTCAGAATCCTCTTCTGAATC).

#### jip-1a/dE1-19 knock-out

*eft-3p::Cas9::U6p::jip-1p sgRNA2* (GCCGACGGTTATTCGGAGGGT)

*eft-3p::Cas9::U6p::jip-1 sgRNA* (GTAACCCAGAGAGGAGATCT)

1749 bp of 5’ homology region

Forward primer: GTGAAATGAGGCGCTGAACTTG

Reverse primer: CGATTTGTGCTGAAATTTTTAAGACTTAAAATTTCAAG

1515 bp of 3’ homology region

Forward primer: TAATTTTTTTTTAATTTTAAAATCAATTTGTCATCC

Reverse primer: CTCCAAGAGTACGCAAACATCTCAC

#### jip-1a/dE12-19 knock-out

*eft-3p::Cas9::U6p::jip-1 E12-19 sgRNA* (GCATCGGAATACGGCGGAGCA)

*eft-3p::Cas9::U6p::jip-1 sgRNA* (GTAACCCAGAGAGGAGATCT)

1709 bp of 5’ homology region

Forward primer: TCTCAGCTGTAACAACAGTTTTGTGTAC

Reverse primer: GGATGGGAGTACCGATTGGAATAC

1515 bp of 3’ homology region

Forward primer: TAATTTTTTTTTAATTTTAAAATCAATTTGTCATCC

Reverse primer: CTCCAAGAGTACGCAAACATCTCAC

### Germline transformation

DNA was injected into the syncytial gonad of adult hermaphrodites (P_0_) as described previously (MELLO AND FIRE 1995). F_1_ progenies expressing the injected fluorescent transgenes were identified and cloned (1 animal per plate), and the F_2_ progenies were screened for transgenic lines.

### Genetic mosaic analysis

Genetic mosaic analysis was performed in animals containing unstable extrachromosomal transgenic arrays that had been passed for at least 6 generations before analysis, as described previously (SAGASTI *et al*. 2001; VANHOVEN *et al*. 2006). The mosaic animals that lose the extrachromosomal transgene in one of the two AWC cells were identified by the loss of the co-injection marker *odr-1p::DsRed* (expressed in both AWC neurons in non-mosaic animals) expression in AWC.

### Mos1-mediated single copy insertion (MosSCI)

Mos1-mediated single-copy insertion of transgenes was performed as described previously (FROKJAER-JENSEN *et al*. 2008). *pAB1.1*::*odr-3p::GFP::jip-1a* (33 ng/μl) or *pAB1.1*::*odr-3p::GFP::jip-1a(vy6)* (45 ng/μl) was injected into *ttTi5605* II; *unc-119(ed3)* III animals. *pCFJ356::odr-3p::tir-1::TagRFP* (67 ng/μl) was injected into *cxTi10816* IV*; unc-119(ed3)* III animals. Each of these transgenes was co-injected with *eft-3p::mos-1* (50 ng/μl), *hsp16.4p::peel-1* (10 ng/μl), *rab-3p::mCherry* (10 ng/μl), *myo-3p::mCherry* (5 ng/μl), and *myo-2p::mCherry* (2.5 ng/μl). The injected animals were grown at 25°C until the worms on the plates were starved. Animals were then heat shocked at 34°C for two hours to induce the expression of the negative selection marker PEEL-1, which kills animals carrying non-integrated transgenes. Animals that moved freely, resulted from the rescue of the *unc-119(ed3)* phenotype, and lost the expression of co-injection mCherry markers were cloned on single plates and verified by PCR to establish single-copy insertion lines.

### Generation of mNG knock-in and knock-out strains using Cas9-triggered homologous recombination

mNG knock-in or knock-out strains were generated using Cas9-triggered homologous recombination as previously described (DICKINSON *et al*. 2013; DICKINSON *et al*. 2015; DICKINSON AND GOLDSTEIN 2016). The sgRNA/Cas9-expressing construct (50 ng/μl) and homology repair template construct (50 ng/μl) were co-injected with three markers, *rab-3p::mCherry* (10 ng/μl), *myo-2p::mCherry* (2.5 ng/μl), and *myo-3p::mCherry* (5 ng/μl), into N2 animals cultured at 20 °C. The repair template construct’s SEC cassette, flanked by LoxP sites, contains transcriptional terminators, a dominant-roller phenotype marker *sqt-1(e1350),* Cre driven by a heat-shock promoter, and a hygromycin resistance gene. Injected worms were allowed to lay eggs at 25 °C. Hygromycin (Gold Biotechnology) was applied directly to the plates at a final concentration of 250 μg/ml three days later, and the plates were cultured at 25 °C. The animals that showed a roller phenotype and lost red fluorescent co-injection markers were cloned onto single plates three to four days later to establish insertion lines. L1 progenies of the insertion lines were heat shocked at 34°C for four hours to induce Cre expression, then cultured at 20 °C for 5-7 days. The F_2_ progeny of heat-shocked worms that moved freely (non-rollers) and expressed mNG were cloned to establish SEC-excised lines. The correct insertion of the mNG marker and the correct excision of SEC were verified by PCR and sequencing.

### Live imaging of transgenic animals expressing fluorescent proteins

Animals, anesthetized with 5 mM sodium azide (Sigma) or 7.5 mM levamisole (Sigma), were mounted on 2% agarose pads on microscope slides. Images were obtained using a Zeiss Axio Imager M2 microscope, equipped with a motorized focus drive, a Zeiss objective EC Plan-Neofluar 40x/1.30 Oil DIC M27, a Piston GFP bandpass filter set (41025, Chroma Technology), a TRITC filter set (41002c, Chroma Technology), a Zeiss AxioCam 506 mono CCD digital camera or a Hamamatsu digital camera C11440, and a Zeiss Apotome system. Images were captured using Zeiss ZEN imaging software.

### Quantification of fluorescence intensity

Animals from each set of experiments were imaged at L1 with the same exposure time. Fluorescence intensity was measured using ImageJ or Zeiss ZEN. In Figures 2D, S1A, and S1B, GFP fluorescence intensity in the AWC axon and cell body was measured in the maximum intensity projection of Z-stack images. In Figures 3B, 3D, 3F, and S1D, a single focal plane with the brightest fluorescence in the AWC cell body was selected from a stack of images to compare fluorescence intensity. mNG and GFP fluorescence intensity were normalized by the TagRFP intensity measured in the same AWC cell in Figures 3B and 3D, respectively.

### Time-lapse imaging of protein trafficking

Worms in the L1 larval stage were anesthetized with 7.5 mM tetramisole (Sigma) and mounted onto 2% agarose pads on microscope slides for imaging. Our previous study showed that 7.5 mM tetramisole, compared to 0.5 mM, 1 mM, and 2 mM, did not significantly affect the axonal transport of TIR-1::GFP along AWC axons (SIETE *et al*. 2024). Time-lapse images were acquired for 30 seconds with a speed of 5 (TIR-1::GFP) or 10 (GFP::JIP-1a and GFP::JIP-1a^E58K^) frames per second and an exposure time of 100 (GFP::JIP-1a and GFP::JIP-1a^E58K^) or 200 (TIR-1::GFP) milliseconds using a Zeiss Axio Imager M2 microscope equipped with a Zeiss objective EC Plan-Neofluar 40x/1.30 Oil DIC M27, EC Plan-Neofluar 63x/1.40 Oil DIC M27, a Piston GFP bandpass filter set (41025, Chroma Technology), and a Hamamatsu digital camera C11440. Acquired images were analyzed to generate kymographs using Fiji (SCHINDELIN *et al*. 2012) with the KymographClear macro (version 2.0a) (MANGEOL *et al*. 2016). The percentage and velocity of moving events were measured using KymographDirect (version 2.1) (MANGEOL *et al*. 2016).

---

**Figure S1.**
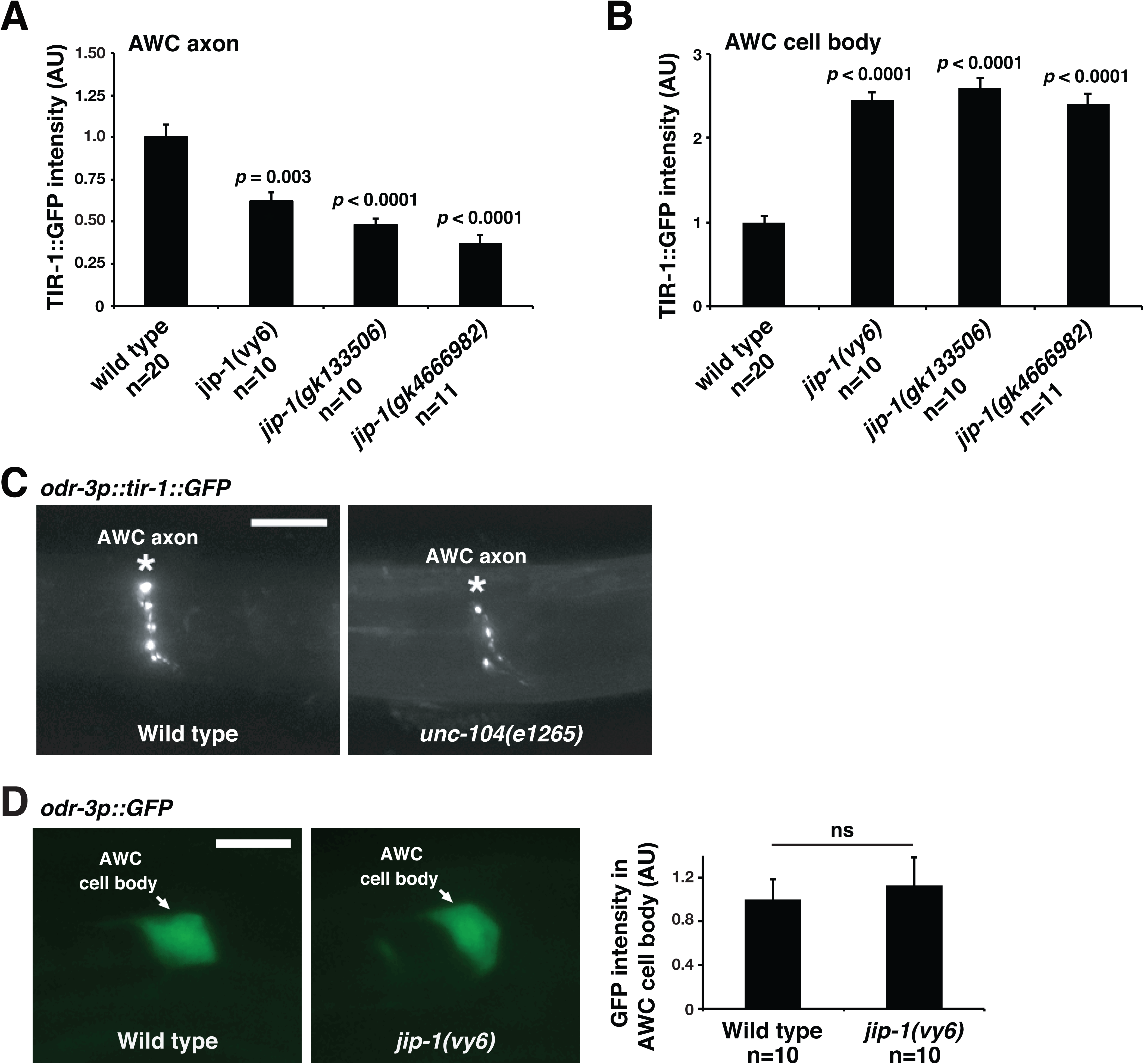
Expression of TIR-1::GFP in wild type*, jip-1* mutants, and *unc-104* mutants. (**A, B**) Quantification of TIR-1::GFP fluorescence intensity, expressed from a stably integrated transgene *odr-3p::tir-1::GFP*, in the AWC axon (A) and cell body (B) at the L1 stage. AU, arbitrary unit. n, the total number of animals analyzed. Student’s *t*-test was used for statistical analysis. Error bars, standard errors of the mean. (**C)** Images of wild type and *unc-104(e1265)* mutants expressing TIR-1::GFP in AWC cells from a stably integrated transgene *odr-3p::tir-1::GFP* in L1. The anterior is left, and the ventral is down. Scale bar, 5 μm. (**D**) Left panels: Images of wild type and *jip-1(vy6)* mutants expressing *odr-3p::GFP* in AWC cells at the L1 stage. The anterior is left, and the ventral is down. Scale bar, 5 μm. Right panel: Quantification of GFP fluorescence intensity in the AWC cell body. In each animal, fluorescence intensity was quantified from the single focal plane with the brightest GFP expression in the AWC cell body. AU, arbitrary unit. n, the total number of animals analyzed. Student’s *t*-test was used for statistical analysis. ns, not significant. Error bars, standard errors of the mean.

**Figure S2.**
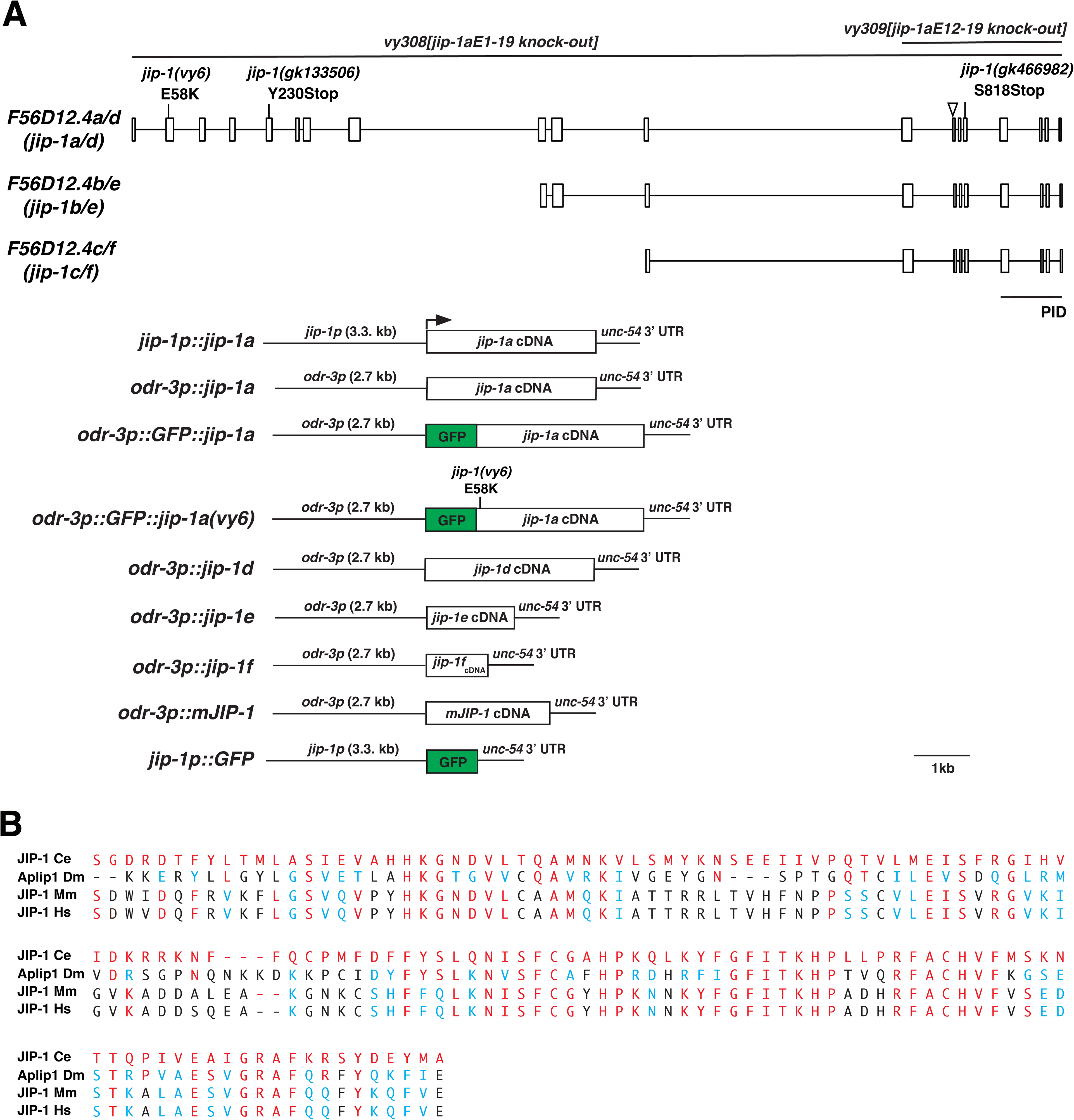
Gene structure of *jip-1*. (**A**) Top: Genomic DNA structure of *jip-1* isoforms. PID, phosphotyrosine-interaction domain. The arrowhead indicates the position at which the amino acid sequence differs between *C. elegans* JIP-1 protein isoforms a/e/f (SFFSPD) and b/c/d (Y). Bottom: Structure of transgenes used to rescue *jip-1(vy6)* mutants and the analysis of *jip-1* expression pattern. (**B**) Amino acid sequence alignment of PID domain in *Caenorhabditis elegans* (Ce) JIP-1 with homologs in *Drosophila melanogaster* (Dm) (PELLET *et al*. 2000), *Mus musculus* (Mm), and *Homo Sapiens* (Hs). The alignment was performed using UniProt (https://www.uniprot.org/align/). Red, identical; blue, similar; black, not similar.

**Figure S3.**
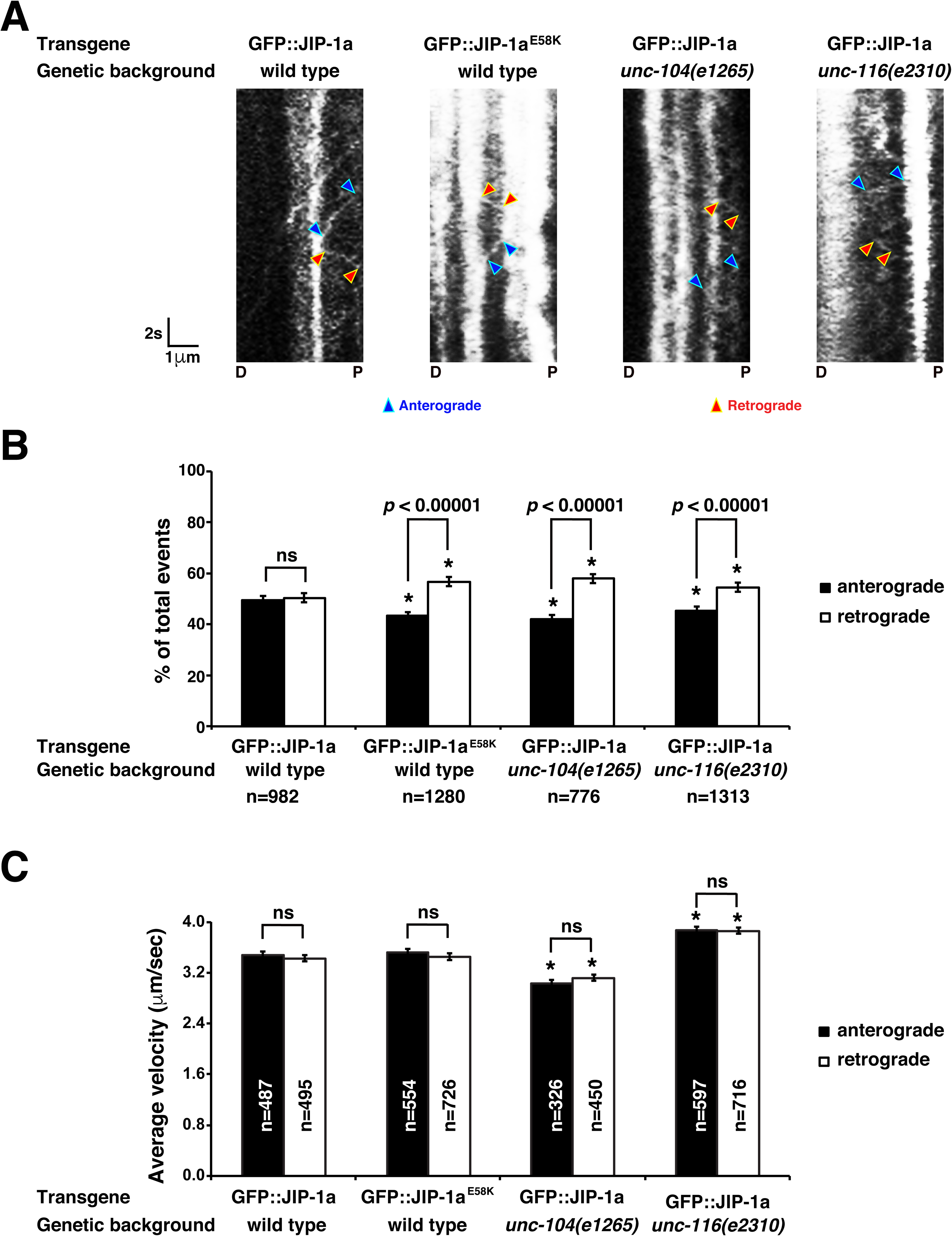
UNC-104 and UNC-116 are required for the dynamic trafficking of JIP-1 in the AWC axon. (**A**) Representative kymographs of GFP::JIP-1a and GFP::JIP-1a^E58K^ movement in the AWC axon at the L1 stage. (**B, C**) Quantification of the percentage of anterograde and retrograde GFP::JIP-1a and GFP::JIP-1a^E58K^ trafficking events (B) and the average velocity of trafficking events (C). Asterisks indicate significant differences (*p* < 0.0001) in the same direction of movement between GFP::JIP-1a versus GFP::JIP-1a^E58K^ or GFP::JIP-1a in wild type versus *unc-104(e1265)* or *unc-116(e2310)* mutants. *P*-values were determined by a *Z*-test (B) or Student’s *t*-test (C). Error bars represent the standard error of the proportion (B) or the standard error of the mean (C). ns, not significant. n, number of trafficking events.

## Notes

### Competing Interest Statement

The authors have declared no competing interest.

